# Encoded and non-genetic alternative protein variants expand human functional proteome

**DOI:** 10.1101/2025.02.17.638604

**Authors:** Vyacheslav Tretyachenko, Tehila Leiman, David Morgenstern, Yishai Levin, Omer Asraf, Orna Dahan, Dvir Dahary, Yitzhak Pilpel

**Affiliations:** Department of Molecular Genetics, Weizmann Institute of Science, Rehovot, 76100, Israel; Nancy and Stephen Grand Israel National Center for Personalized Medicine G-INCPM, Weizmann Institute of Science, 760001 Rehovot, Israel

## Abstract

Each stage of the Central Dogma contributes to proteome diversity through mechanisms such as heterozygosity, somatic mutations, transcriptional errors, and translational errors. As a result, a diverse array of protein variants can coexist within a single proteome, such as that of humans. However, until now, methods to detect, quantify, and evaluate the functional consequences of these variants have been lacking. Here we examined a large-scale proteogenomic dataset from 29 healthy human tissues and uncovered 13,910 confidently localized variants representing 7,215 unique single amino acid substitutions co-existing alongside their corresponding reference proteoforms. We found that the abundance of both genetic (SNP’s, somatic mutations) and mistranslated protein variants mirrors their allele frequencies in the human population. Moreover, we show that non-genetic substitutions may provide a distinct route for exploring protein sequence space, circumventing the mutational constraints imposed by the genetic code. In addition, we provide experimental validation of non-genetic substitution on selected purified proteins. We demonstrate specific and recurring non-genetic variation patterns upon amino acid starvation in proteome-wide analyses of cancer-derived cell lines and identify hundreds of substituted non-genetic proteoforms that recur consistently in multiple healthy individuals or map to annotated protein functional sites. We propose that these substitutions constitute a novel class of functional protein phenotypic variants. Collectively, our findings indicate that non-genetic amino acid substitutions in human proteins provide an abundant source to expanding the functional proteome.

## Introduction

The human genome provides the blueprint for the majority of proteins. However, it has become clear that the complexity of the human proteome extends well beyond the canonical set of approximately 20,000 protein-coding genes^1^. The molecular determinants of cellular phenotype arise from an intricate interplay of differentially spliced transcripts^2^, posttranslational modifications^3^, heterozygotic allele variants^4^, somatic mutations^5^, and additional phenotypic variants generated by transcriptional or translational errors^6,7^. Translation errors can provide diversity to the proteome by multiple means, primarily by featuring amino acid substitutions^7–9^, ribosomal frame-shifting^10^ and STOP codon readthrough^11,12^. Together, these diverse processes can create an enormous range of proteoforms, offering myriad ways to fine-tune the cellular phenotype.

The diploid nature of the human genome provides a distinct mechanism for increasing protein phenotypic heterogeneity by expressing both maternal and paternal heterozygotic variants of genes. These alternative proteins may differ subtly in structure, function, stability, or solubility, resulting in various positive and negative effects on cellular fitness^13^. Notably, co-expression of alternative allelic variants can further expand the proteome’s diversity when combined with allele-specific gene regulation^14,15^. For instance, genes essential for cell differentiation or metabolism often display differential expression of maternal and paternal alleles, leading to subtle functional distinctions in protein isoforms^4,16,17^. Heterozygosity can enhance resilience against pathogenic stressors^18,19^, mitigate the impact of deleterious homozygous mutations^20^, or influence therapeutic responses^21^. Altogether, the interplay of maternal and paternal allelic contributions exemplifies the innate complexity and adaptability of the human proteome.

Somatic mutations that occur post-fertilization, and that are not inherited through the germline to the next generation, also add to the diversity of proteins encoded in an organ and tissues in the body. Yet only rarely manifestation of somatic mutation at the proteome level of tissues has been documented. Although somatic mutations are frequently linked to senescence, malignant cell growth and evolutionary pressures within tumors, they can also provide selective advantages^22,23^. The foremost example is somatic mutagenesis followed by error-prone DNA repair in B-cells, which deliberately fosters diversity in immunoglobulin production^24^, yet demonstration of the DNA variants at the protein level, and their quantification is still limited^25^. Beyond this well-known immunological strategy, other instances of beneficial somatic mutations, such as revertant mosaicism in genetic disorders, have been reported, though they remain relatively scarce and primarily confined to specific clinical contexts^26,27^. Still, as somatic variation is omnipresent, pervasive, and unique to each individual, unraveling its broader role in human health and adaptation represents a considerable challenge^5^.

In addition to genetically encoded alternative proteoforms, protein variants resulting from errors in transcription and/or translation (phenotypic variants) may represent a large fraction of the heterogeneous proteome. Phenotypic variants are orders of magnitude more prevalent than DNA mutations, ranging between 10^-5^ and 10^-2^ per RNA and protein residues, respectively^28^. Originally mapped systematically in microbes^7,8^, recent analyses were expanded to flies and human^9,29^ revealing a codon usage effects and amino acid levels in cells as key determinants of translation infidelity. With ∼2×10^9^ protein molecules in a human cell, and ∼10^12^ amino acids in them, an averaged translation error rate of 10^-3^ amounts to 10^9^ mistakenly incorporated amino acids, occurring in the cell’s proteome and many of the proteins bearing phenotypic mutations to some extent^30^. A prime characteristic of phenotypic mutations is that they cast a significant burden on organisms. Proteins translated with errors tend to misfold, aggregate, engage in spurious interactions, saturate the protein quality control machinery^31–33^, and cause human diseases^34^, e.g. neurodegeneration^35,36^. In parallel, an increasing number of reports now indicate that some errors in translation can be advantageous at the cellular level ^31,37–39^. In cancer, a recent summary provides evidence that translational errors contribute to proteome diversity and population heterogeneity, thus contributing to the tumor’s capacity to evolve^40^. On an evolutionary scale, phenotypic errors might open evolutionary paths otherwise precluded by epistatic interactions^41^, to facilitate the purging of deleterious mutations^42^, or expose new hidden functionalities^43–45^. Whether deleterious or beneficial, the rates and nature of phenotypic mutations appear to be regulated^37^ and the tendency to make or avoid them is postulated to be genetically encoded in the DNA^7,46–48^.

In this work, we present a unique perspective on alternative proteoforms in human cells and their potential functional implications. We developed a novel proteogenomic pipeline for the identification of single amino acid variants and applied it to the datasets of (i) healthy human tissues, (ii) amino acid starved cancer-derived cell lines, (iii) proteasome-inhibited U2OS cells and (iv) enriched/purified human proteins ^9,49–52^. Our analysis explored and quantified thousands of unique genetically encoded and non-genetic protein variants. We found that the abundance of these substitutions correlates negatively with the predicted pathogenicity of the resulting variant and the variant’s allele frequency in the human population. We observe a general tendency of non-encoded substitutions to introduce amino acids with distinct biochemical properties and beyond the single-base mutation range of their codon (*i.e.* non-cognate substitutions).

## Results

### Substitution search and validation pipeline

We have developed an improved approach towards detection of amino acid substitutions within proteins, using mass spectrometric (MS) proteomic data. A previous method is based on the MaxQuant dependent peptide search algorithm^7,53^ that was originally developed for mapping post-translational modifications. That method became adopted for proteome-wide amino acid substitution detection^7,29^. However, the drawback of the dependent peptide search for single amino acid substitution detection is in its low substitution detection rate. This results from its broad purpose – identification of as many modified peptides as possible. Yet, the breadth of the search is not beneficial when one is interested in a specific variant category such as single amino acid substitutions^54^ Here we present a novel amino acid substitution detection pipeline which improves upon the previous methodology and builds on recent advances in mass spectrometry data analysis^55–57^. Our pipeline utilizes (i) published machine-learning assisted re-scoring of peptide-spectrum matches (PSM’s)^55^, (ii) 2-stage false discovery rate (FDR) error controlled proteomic search to separate search spaces of the reference/substituted peptide classes^57,58^ and (iii) published peptide-centric multistage PSM validation procedure and mass shift localization recently introduced in PepQuery2 and pyAscore tools^59,60^.

In short, our pipeline consists of three search steps, detailed below and summarized in Fig. 1a:

1) Raw mass spectrometry data is searched with a reference proteome database, and unidentified spectra are aggregated into the new raw files.
2) Identified reference peptides are used to generate a new sequence database of candidate substituted peptides. Identified reference peptides are *in silico* mutagenized - each amino acid position of the reference peptide is systematically exchanged for every other amino acid, resulting in distinct substituted peptide candidates. Unidentified spectra from step (1) are searched with MSFragger^61^ for the substituted peptide candidates and quantified with IonQuant^62^.
3) Every PSM of the identified substituted peptide candidate is further validated with peptide-centric search engine PepQuery2 and substitution position is confidently localized by Ascore statistical framework^56,59,60^. PepQuery2 validation consists of competitive scoring of the tested peptide match against (i) 10,000 of its own shuffled variants, (ii) peptides from the reference proteome, (iii) post-translationally modified (PTM’s) reference peptides (including labile modifications) and (iv) single amino acid variants of other peptides from the reference proteome. A candidate peptide and its spectrum are considered valid only if the candidate peptide score is higher than any of the tested competing variants (i)-(iv) and the putative amino-acid substitution mass shift is confidently localized to the expected reference residue. PepQuery does not rely on the traditional target-decoy FDR estimation strategy, hence it provides statistically independent way to validate previously identified substituted peptide candidates with experimentally estimated FDR of 2%^56^.

**Figure 1.**
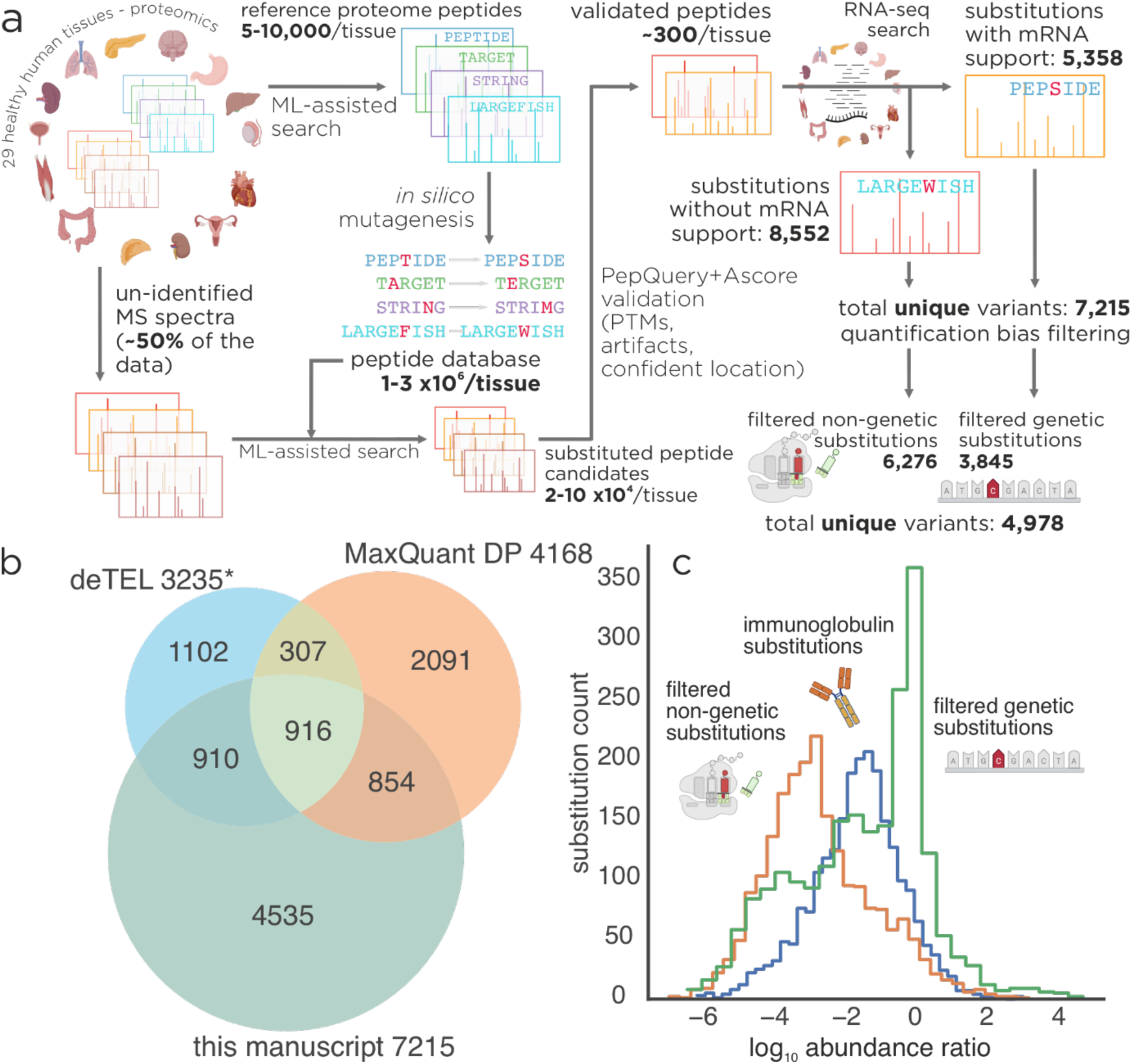
Substitution strategy overview and outputs. (A) Overview of the amino acid substitution detection pipeline utilized in this work. Raw proteomic data are split into identified and un-identified subsets, identified peptides are *in silico* mutagenized and searched for within the un-identified subset of the raw data. Substituted peptide candidates are validated by PepQuery2, localized by Ascore, verified for RNA-seq evidence in transcriptomics data and further filtered for the quantification purposes. (B) Substituted peptides identified by MaxQuant dependent-peptide search protocol^7^ (orange), open-search based deTEL pipeline^8,64^ (blue) and the method presented here (green) from the same dataset. (C) Relative abundances of substituted peptides and their reference variants as ratios of their precursor (MS1) intensities. Blue – substitutions without mRNA-seq support, orange – substitutions found in immunoglobulins (with or without mRNA-seq support), green – substitutions with mRNA-seq evidence

To further validate our variant peptide identifications, we compared experimental and predicted MS2 spectra already used as a feature in Percolator rescoring framework within the MSFragger search. We observed strong agreement between experimental and theoretical values (median spectral similarity > 0.9; Extended Data Fig. 1, sample mirror spectra Extended Data Fig. 2 and Supplementary Fig. 1-7). Further, predicted and experimental retention times (also utilized in MSFragger search) for substituted precursors were highly correlated (Pearson’s r = 0.91, P < 1 × 10⁻¹⁶; Extended Data Fig. 1). Predictions agreed less well with experimental spectra and retention times for substituted peptides than for their reference counterparts (median spectral similarity 0.91 vs 0.96, Pearson correlation of retention times 0.91 vs 0.94 in substituted and reference peptides respectively, Extended Data Figure 1 and Supplementary Fig. 8), consistent with (i) the absence of these peptidoforms from prediction-model training sets and (ii) published observations on drop in prediction performance for low-abundance peptides with sparse fragment-ion coverage. Focusing on confidently identified reference proteome precursors we independently confirmed that agreement with predicted fragmentation spectra indeed decreased with abundance. (Supplementary Fig. 8) ^55,63^. Together, these orthogonal validations strengthen confidence in correct precursor identification.

Identified and validated substituted peptides are then backtracked to sample-specific RNA-seq data to determine whether the observed substitution originated from a mutation in DNA/mRNA or, in case of only reference variant observed in transcriptomic data, by non-genetic mechanism such as ribosomal mistranslation. For analyses where misclassification of a non-genetic substitution as a genetic variant could materially bias the results (e.g., comparisons to allele frequency or AlphaMissense pathogenicity), we used a more conservative subset requiring ≥10 sequencing reads supporting the reference allele and 0 reads supporting the alternative allele.

For analyses aimed at describing aggregate properties of substitutions (such as overall substitution spectra, mutational distances, or global abundance trends) this additional stringency is less critical, and we therefore used the broader set defined by ≥1 reference read and 0 alternative reads.

We compared our pipeline with the previous method based on MaxQuant dependent peptide search as well as more recent MSFragger’s open-search based protocol deTEL^7,64^. The substitution detection identification was performed as described in corresponding publications and identified peptides from all three pipelines were validated by PepQuery2^7,8,59^.

In comparison to deTEL and MaxQuant dependent peptide search, we were able to identify, validate and localize >60% and >40% unique substituted peptides on highly fractionated human proteome data (Fig. 1b)^49^. Throughout this work, unique substitutions refer to the number of distinct substituted peptide sequences identified, whereas total substitutions refer to the cumulative count of their observations across all tissue samples (i.e., the same substituted peptide identified in n samples contributes 1 to the unique count and n to the total count). For comparability, we report only MaxQuant- and deTEL-derived substitutions that were independently validated by PepQuery2 using the same settings as in our pipeline. Using MaxQuant dependent peptide search we detected 4,168 unique substituted peptides across all human tissues. (Fig 1b, red). Out of these, 1,770 (42%) were also detected by our pipeline (Fig 1b, red/green circle overlap). Using deTEL, we identified 3,235 unique substitutions (Fig. 1b, blue) and 1,826 (56%) were also detected with our pipeline (Fig. 1b, blue/green circle overlap). Using all three amino acid substitution pipelines yields 33% more variants than reported using our methodology (Fig. 1b, non-overlapping parts of red/blue circles). Substitutions detected by our method, MaxQuant dependent peptides search and by deTEL are reported in Supplementary Table 1, 2 and 3 respectively. In this work, we chose to characterize the data identified and quantified by a single approach presented here.

In the initial MSFragger search of our pipeline we identified 915,484 candidate substituted peptide spectra. This count is inherently inflated by false positives due to the large database size and shortcomings of the FDR estimation by target target/decoy competition as often witnessed in metaproteomic studies^58^. Hence, we evaluated all PSM’s corresponding to these candidates with PepQuery2 which filtered out 93% of the candidate PSMs. To further improve the substitution identification confidence, we required that (i) substituted positions were covered by at least two fragments and we (ii) excluded all substitutions on peptides N’ terminus due to low detectability of the corresponding b1 ions^65^. This yielded an intermediate dataset of 76,516 candidate peptides with single amino acid substitution. As a final validation step, we applied two additional filters. First, we removed spectra that exhibited a better MS2 match to the corresponding reference-proteome peptide when precursor mass was ignored, thereby excluding potential reference variants carrying labile modifications. Second, for substitutions lacking genetic support, we excluded spectra with ambiguous substitution site localization as determined by the Ascore algorithm ^60^. Ascore localizes a modification by scoring how well the observed MS/MS spectrum contains the site-determining fragment ions uniquely predicted for that mass-shift (using a binomial model for matching peaks) and reports the best site and a confidence score based on the score difference to alternative sites. We only considered identification where the highest scoring site coincided with the substituted position and its score was higher than for any other localization within the peptide sequence. This localization filter retained only confidently assigned substitutions, ensuring that the observed precursor mass shift could not be attributed to a modification elsewhere in the peptide sequence. We did not apply strict localization filtering when the substitution was independently supported by mRNA-seq data. After filtering, we obtained 13,910 validated peptides with confidently localized substitution sites in non-genetic variants (Supplementary Table 1).

For each detected substitution we assigned an abundance ratio defined as ratio of precursor (MS1) intensities of the substituted and reference peptide. We observed a wide range of substitution abundance ratios from 10^-7^ up to 10^2^ (Fig. 1c). In addition, we analyzed transcriptomic data provided with the original dataset^49^ to determine potential genetic or non-genetic origin of the observed substitutions.

### Amino acid substitutions are prevalent within the human proteome

We analyzed mass spectra from the healthy human tissues for which RNA-sequencing data were available and detected 13,910 amino-acid substitutions complying with our strict filtering criteria^49^. Out of these, 7,215 represented unique substitutions. As a result of amino acid substitution-focus, we achieved at least 40% increase in number of detected substitutions in comparison to previously reported methods, when applied to the same data (Supplementary Tables 1, 2 and 3)^7^. During the analysis we have discovered a systematic bias towards increased detection rate of amino acid variants in the vicinity of the N’ terminus of tryptic peptides. This bias was also observed in the ”ground truth” substitution group, - variants supported by RNA sequencing and expected to be observed in proteins, indicating that it does not represent false positive identifications within our data and compromise the overall statistics of amino acid substitution analysis (Extended Data Fig. 3, left). Nevertheless, to avoid potential quantification bias associated with trypsin cleavage specificities upon the changing substrate (substituted protein)^66^ in the vicinity the cleavage site, we only considered substitutions that fulfill the following criteria: (i) found on peptides derived from complete or near-complete (one missed cleavage maximum) protein digestion by trypsin, (ii) within a distance of at least three amino acid positions away from either trypsin cleavage sites on N’ and C’ peptide termini and (iii) not originating, nor leading from/to lysine or arginine residues, the two amino acids associated with trypsin activity. We refer to this dataset as the ”filtered data”, which we use only for analyses requiring MS1 abundance quantification (Extended Data Fig.3, right). This filtering retained 10,121 substitutions (out of which 5,257 are unique) (Supplementary Table 4). In cases where we only evaluated whether a substitution is present or not, without considering quantification of the alternative protein abundance, we employed the previously described ”expanded set” of 13,910 substitutions (Supplementary Table 1).

Our collection of detected and filtered substitutions contains both genetically encoded amino acid variants (due to either somatic mutations or heterozygotic allele expression) and phenotypic variants of proteins resulting from errors in transcription and translation. We utilized the transcriptomic sequencing data accompanied with the proteomic dataset to further dissect the origins of the observed amino acid substitutions and determined if they are likely derived from DNA substitution, or phenotypically by transcription/translation errors. For this purpose, we assigned an RNA read coverage ratio for each MS1-derived substitution intensity. In analogy with MS substitution abundance ratio, RNA coverage ratio represents the log_10_ ratio of read counts that encode the observed amino acid substitution to the counts of the reads that encode the reference amino acid. This ratio represents a direct measure of a genetic variability on mRNA level and an indirect estimation of it on the level of DNA assuming the low frequency of transcription errors. In many cases, RNA coverage ratios suggest only small fraction of substituted transcript in the sample which raises a challenge in distinguishing between biological origin of substitution and mere technical artefact, likely mutations originated from transcriptomic library preparation. For each of the substitutions with RNA-seq evidence we calculated the binomial probability that *k* observations of a substitution within its *N* sequencing reads are not of technical origin assuming the error rate of transcriptomics pipeline 0.1%^67^. The rationale for this step is to re-evaluate the genetic origin for the substitutions supported by relatively few reads considering the sequencing depth on this position. We define the MS-observed substitution to have RNA evidence if the FDR-corrected probability of technical origin is lower than 0.05. This filtering yielded 5,358 RNA-Seq supported substitutions with high confidence of non-technical origin (Supplementary Table 1).

When the MS and RNA-seq derived substitution ratios are plotted against one another, several substitution groups can be observed (Fig. 2). The first group is centered around 0-0 coordinates, where both proteome MS and RNA-seq abundance ratios of the reference amino acid and the substituted one are about equal (log ratio around 0). Equality of the RNA read counts for the reference and substituted variant suggests heterozygosity of the individual sample donor, and about equal mRNA expression level of the two alleles. Interestingly the deviations from equal MS substitution abundance despite the roughly equal RNA coverage ratio may represent cases of higher or lower alternative proteoform abundance in comparison to the reference. These cases can exemplify either (i) differentially translated allelic variants or (ii) differential stability of the proteins with the reference or alternative forms. We will discuss outstanding manifestations of substituted protein pathogenicity/stability in more detail in the following sections.

**Figure 2.**
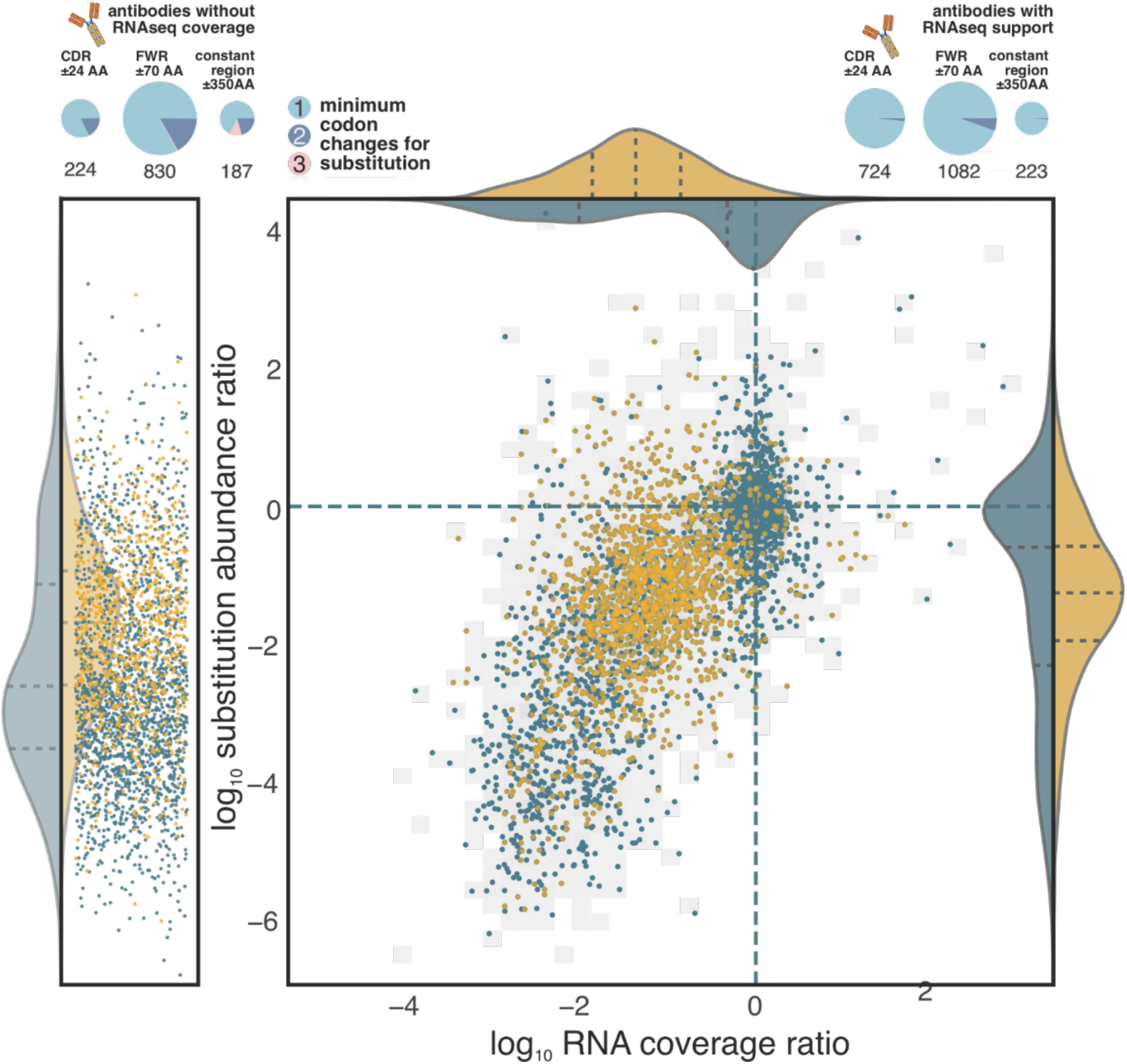
Quantification of substituted variants. Comparison of substitution abundances provided by RNA-seq coverage and MS1 intensity ratios of substituted and reference peptides. Right – substitutions found on both protein and mRNA level and their corresponding abundances in comparison to the reference peptide and transcript. Grey background area - position of data present in expanded set but filtered out in the filtered set. Left – MS observed substitutions without evidence in RNA-seq data and sequencing coverage ≥ 1 supporting only the reference protein variant. Yellow – members of the immunoglobulin superfamily, blue – substitutions of all other (non-immunoglobulin) proteins. Violin plots on x/y axes represent distributions of immunoglobulin (yellow) and non-immunoglobulin (blue) coverage (x-axis) and abundance ratios (y-axis). (Top) immunoglobulin substitutions, split by whether a supporting genetic variant was detected (right) or not (left). Pies show substitution counts (below) and proportions (size) across CDR, framework (FWR), and constant regions (left to right). Colors indicate substitutions reachable by one (light blue), two (dark blue), or three (pink) codon mutations.

Next section of the abundance plot consists of the area spanning the values of -1.5 to -3 (substitution abundance ratios ranging from 1:10 to 1:1000) in both proteome MS, and RNA-seq coverage ratio plot. These substitutions are deduced to represent genetically encoded, low-abundance variants in the dataset. Although these variants could represent transcriptional errors, we suggest that these substitutions outline somatic variation as transcription infidelity was observed at much lower RNA coverage ratios (one nucleotide per 10^4^-10^5^ transcription events)^6^. Notably, ∼50% (630 out 1,238) of the genetically encoded substitutions in the somatic variant group were members of the immunoglobulin superfamily. Substitutions of immunoglobulins are illuminated by a unique color on the plot, Fig. 2. As immunoglobulins circulate systemically: a peptide identified by MS in one tissue need not have been translated there, we cannot disentangle secreted antibody variants from genuine local mistranslation events. We flag immunoglobulin detections in Supplementary Tables 1 and 4 and summarize their substitution trends in Figure 2 (top).

The leftmost section of the plot represents all MS observed substitutions which were not supported by RNA sequencing data (Fig 2, left). This cluster represents 60% of the reported amino acid substitutions. Variants in this region could be generated either by (i) RNA-seq coverage limitations, which could have resulted in omission of reads that cover the substituted peptide sequence even though it exists, (ii) unsampled somatic variants from different areas of the tissue, or (iii) mistranslation errors during proteins synthesis. Substitutions found in this region of the abundance ratio plot had at least 1 RNA read that confirms the reference amino acid but lacked RNA evidence for the alternative variant identified by mass spectrometry (Fig. 2, left). When we employ a higher threshold on RNA coverage ≥10 reads mapped to the reference variant, we still detect 6,510 (76%) of total 8,552 substitutions without RNA-seq support for the substituted protein variant. Although we cannot exclude the genetic origin of the detected substitutions completely, we assume that the majority are errors in translation or transcription as the frequency of these phenotypic processes were evidenced to be 2-3 orders of magnitude higher than the DNA polymerase mutation rate ^6–8,68,69^. Indeed, the average log abundance ratio of these substitutions averages to -2.6 (∼1/500, Fig. 1c) similar to the values reported in other high-throughput analyses of translational fidelity ^7,8,70^.

Intriguingly, and in agreement with recent investigations of human proteomics data^70^, 539 substitutions, that make up 2.3% of the 13,910 variants in our expanded set, showed an abundance ratio of the alternative to reference variants greater than zero, suggesting that these variants may exceed the abundance of their corresponding reference proteoforms (Supplementary Table 1). Even with stricter requirement for non-genetic substitution origin evidence by a minimum RNA sequencing coverage threshold as high as 100 RNA-seq reads, we identify 68 such non-immunoglobulin derived substitutions lacking genetic support but still exhibiting a log_10_ abundance ratio above zero (Supplementary Table 4). While the idea that non-coded proteins might dominate the proteome is conceptually compelling, we acknowledge the limitations of MS1-based peptide quantification as well as potential confounding factors such as differential peptide ionization efficiency or chromatographic retention that may inflate apparent abundance. Additionally, because most detections arose from low-abundance proteins, quantification of abundance ratios may be biased by reduced MS1 signal-to-noise ratio. Nevertheless, we experimentally assessed ionization efficiencies for seven reference/substituted synthetic peptide pairs selected from cases in which the substituted peptide exhibited higher MS1 intensity than its reference counterpart. In four out of seven pairs, the substituted variants showed higher ionization efficiency than the corresponding reference peptides (Extended Data Fig. 4). However, the magnitude of these ionization efficiency differences was insufficient to fully account for the observed disparities in MS1 intensities. We therefore propose that a combination of factors (including differential ionization efficiency, noisy quantification at low signal levels, and chromatographic biases) may overestimate the abundance of substituted (non-genetic) forms by up to 100–1,000-fold relative to their reference variants. Accordingly, while the apparent abundance of substituted variants may be inflated by technical factors, their true levels are likely more modest. Importantly, they could still reach biologically meaningful abundances (on the order of single-digit percentages relative to the reference form) consistent with our ability to confidently detect and validate such variants even in low-abundance proteins.

### Expression ratios of genetically encoded protein variants recapitulate their human population frequency

We analyzed the abundance ratios of genetically encoded substitutions (heterozygous alleles and somatic mutants) in light of their population frequencies as found in the Genome Aggregation Database gnomAD 4.1 database^71^ (Supplementary Table 5). We began our analysis by examination of heterozygotic protein variants. To classify identified substituent as heterozygotic allelic variant we required its log_10_ mRNA coverage ratio between –0.33 and 0.33 (corresponding to ∼twofold differences in their transcript abundances^72^) and minimal sequencing coverage of 10. In total 630 unique variants fit into these criteria and for 498 of them allele frequency annotations are available in the gnomAD database. The majority of these allelic variants expectedly clustered around the 0 region of the y-axis (i.e. about equal protein abundances of reference and substituted variant) of the plot (Fig. 2). Nonetheless, some variants showed markedly different protein abundance ratios than would be expected from their mRNA expression ratios. Remarkably, annotated rare variants in human population also tend to have lower abundance in comparison to the co-expressed reference variants corresponding to the more frequent allele variant. Despite potential confounding effect in differential ionization efficiency of single amino acid variant peptides, we observed a positive correlation between the variant’s population allele frequency and its peptide substitution abundance ratio (Pearson 0.24, p-value 8.1×10^-11^, Fig 3a,d). This dependence confirms and broadens recently suggested connection between the translation efficiency of the transcript with a missense mutation and its allele frequency in the population^73^, but it may also result from a lower protein stability of variants that are rare in the human population.

**Figure 3.**
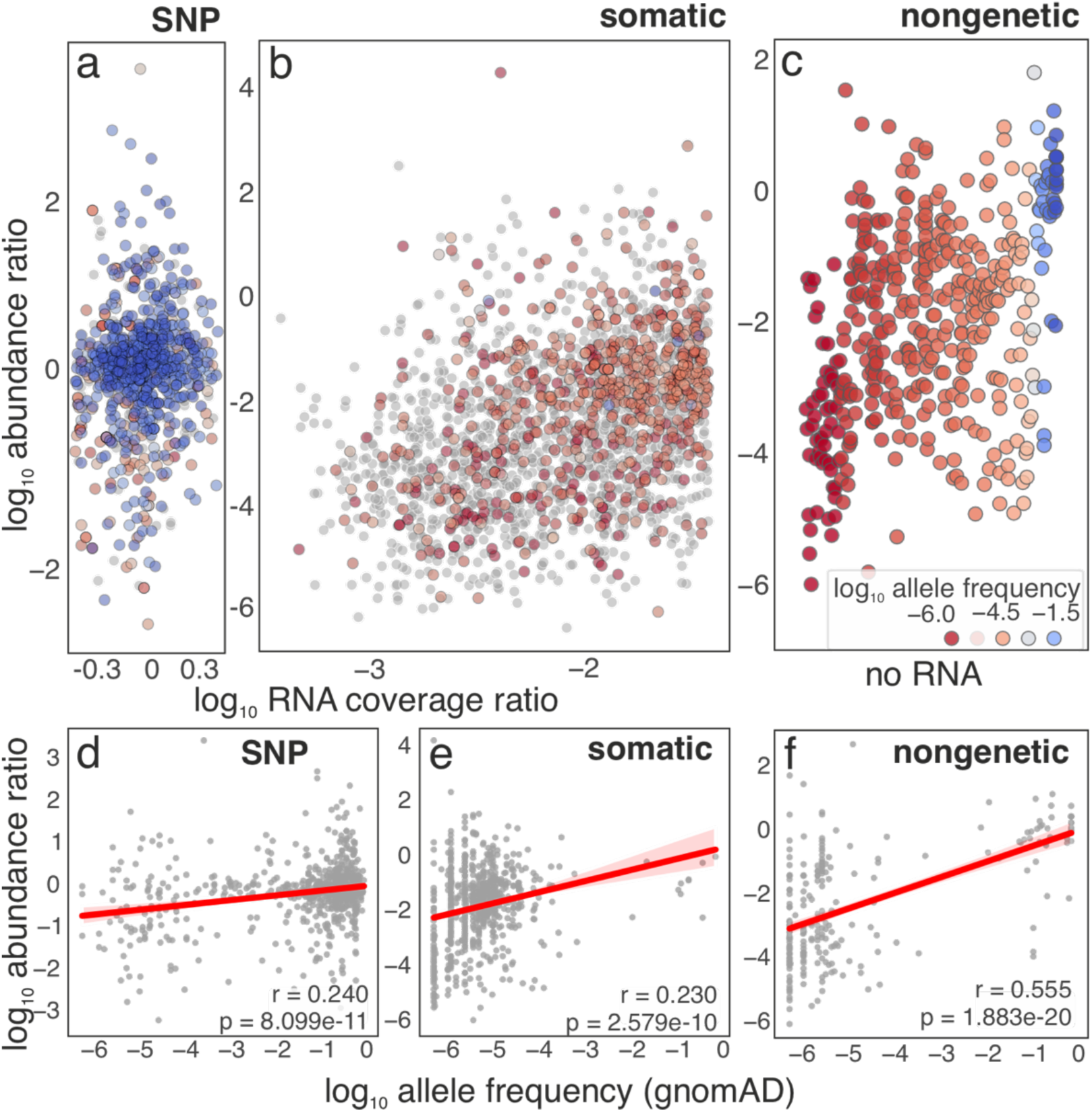
Relation of substitution abundance and its population frequency. (a) Subset of the abundance ratio plot (Fig 2) representing detected heterozygotic allele variants of proteins. RNA coverage ratio (-0.33,0.33) corresponds to ∼twofold difference in the transcript abundance. Colored points represent substitutions also found in human population with the color corresponding to their allele frequency (AF in log_10_ scale, same color scheme as on 3b and 3c). Grey points represent substitutions not found in GnomAD database. (b) Subset of the abundance ratio plot representing somatic variants of the proteins. Colored points represent substitutions also found in human population with the color corresponding to their allele frequency. Grey points represent substitutions not found in GnomAD database. (c) Subset of the abundance ratio plot representing variants without RNA-seq evidence. Sequencing coverage for each substitution is ≥10 reads. Colored points represent substitutions found in human population with the color corresponding to their minor allele frequency. (d) Correlation between population allele frequency and heterozygotic variant substitution abundance ratio. (e) Correlation between allele frequency in population and abundance ratio of somatic variants. (f) Correlation between population allele frequency and protein abundance ratio of variants without RNA-seq support.

Further, we analyzed allele frequencies of the rest of the identified variants with RNA-Seq support, that show a lower substituted-to-reference peptide ratio. We hypothesize that this subset of substitutions located in RNA coverage ratio between -1.5 to -3 represent somatic mutants within the donor tissues. This subset is unlikely to represent heterozygosity because of the large deviation from 1:1 ratio of expression. Using the log_10_ RNA coverage ratio threshold of (-3, -1.5) we identified 868 unique variants in 363 proteins. Notably, 432 variants originated from immunoglobulins (proteins associated with GO terms “immunoglobulin complex” or “antigen binding”), indeed a protein family with extensive somatic mutations^74^. Out of 868 unique substitutions, 363 (including immunoglobulins) were found in the human population as per gnomAD statistics. We observed a similar correlation between the allele frequencies and both ratio of protein abundances as in case of germline variants (Pearson 0.23, p-value 2.58×10^-10^, Fig 3b,e). Our data suggest that the frequencies of *intra-*individual *de novo* mutations recapitulate the same course as the *inter-*individual population allele frequencies established by organismal selection. In other words, these data suggest that cells with somatic mutations that recapitulate more common SNPs in the human population are more likely to expand within the body, potentially due to fitness advantage.

### Non-genetic variants likely consist of mistranslations and show ribosomal error patterns

Variants with RNA-sequencing support for the reference peptide but with no support for the substituted sequence (6,510 substitutions with coverage ≥10, of which 3,561with the coverage ≥100 reads) represent larger part of the reported amino acid substitution cases. As previously discussed, we assume that most of these substitutions consist of translation errors since mistranslation is a major source of cellular phenotypic variation^7,8,31^.

Notably, the abundant substitutions observed in our and other analyses - such as I>V^75^, Y>F^76^ and S>N^77^ reflect high levels of mistranslation within specific substituted positions of proteins rather than a uniform trend across the proteome. Consequently, particular substitutions appear to be more prevalent in some protein contexts than in others, indicating a potential underlying mechanism involving either cis or trans regulation of amino acid substitutions. Our analysis revealed an unexpectedly high prevalence of C>A, C>S, H>D, M>D, and W>D substitutions across the dataset (Fig. 4b). This class of substitutions was abundant among identifications from all three substitution detection pipelines tested in this work (Supplementary Tables 1, 2, and 3). A review of the literature suggests that several of these events can arise from technical chemical transformations: C>A/S has been associated with desulfurization and subsequent rehydration during proteomic sample preparation ^82^; H>D has been reported following long-term storage and oxidation of blood samples ^83^; and M>D and W>D, while described in the chemical literature, has primarily been observed under specific conditions such as transition metal- or ozone-catalyzed oxidation ^84–86^. We therefore consider this group of substitutions to potentially comprise a mixture of non-biological and biological events, as we consistently detect them in additional datasets that were not exposed to overt oxidative damage or desulfurization-promoting reagents and are discussed in the following sections of this manuscript. Accordingly, we flag these substitutions as cautionary in the main text (Fig. 4b) and supplementary data and treat them as non-biological unless supported by orthogonal evidence. All subsequent analyses of non-genetic variants exclude these five substitution types (5,216 variants without RNA-seq support), unless stated otherwise.

**Figure 4.**
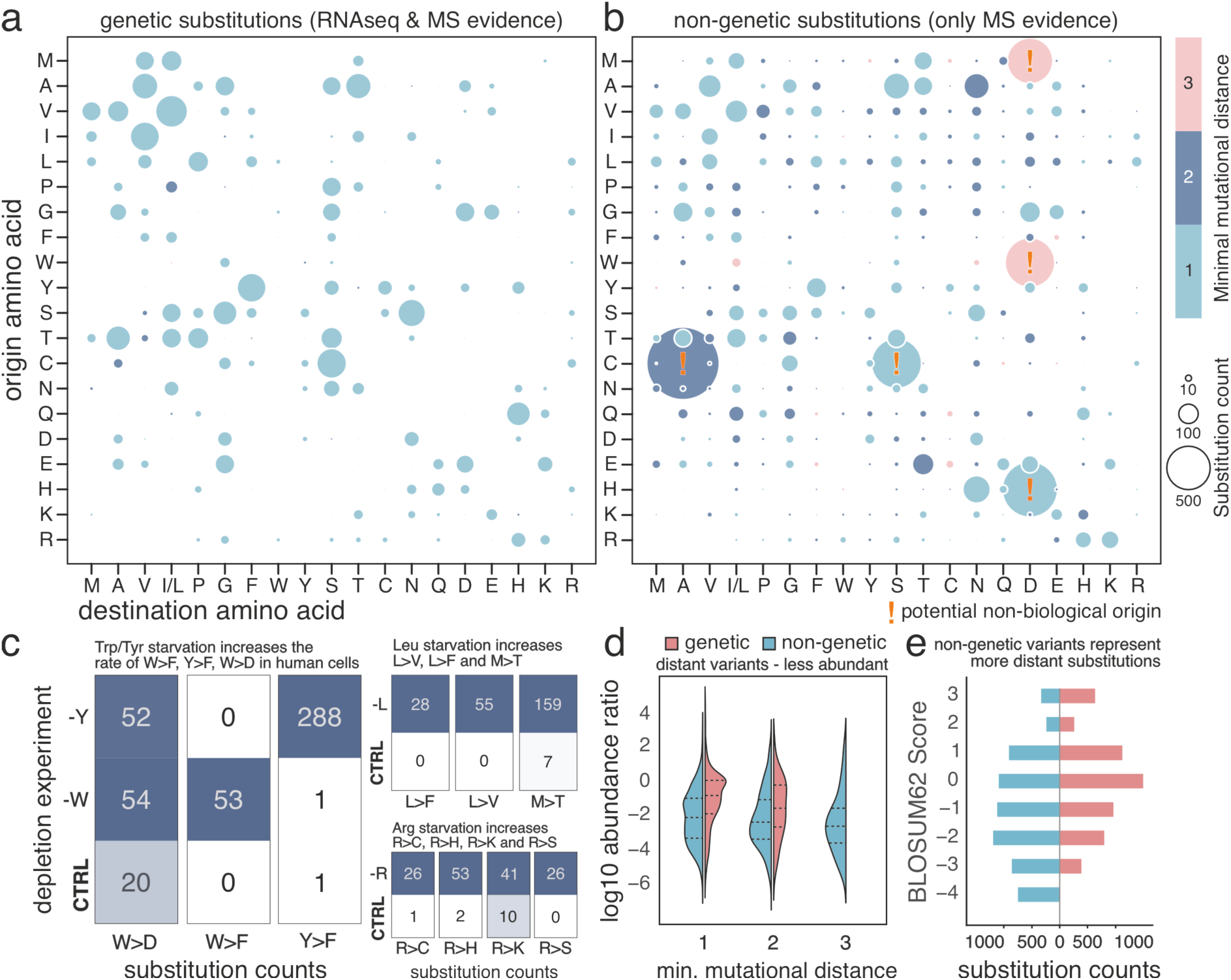
Statistics of amino acid substitutions. (a) Summary of counts of detected genetic substitution types (bubble size) and their minimal mutational distance (bubble color), (b) summary of counts of detected non-genetic substitution types (bubble size) and their minimal mutational distance (bubble color), (c) significant changes in substitution counts between starved and control cells. Left – W/Y starvation, top right – L starvation, bottom right – R starvation, (d) substitution abundance distributions associated with different mutational distances in genetic (red) and non-genetic (blue) substitutions, (e) substitution counts associated with distinct BLOSUM62 scores for genetic (red) and non genetic (blue) substitutions,

The mistranslation spectrum was non-random and directional. The most frequent substitution classes included A>S, A>N, V>I/L, H>N, A>V, E>T, Y>F, A>T, and S>G, indicating that a limited subset of amino-acid changes dominates the observed landscape. Directionality was evident even among relatively conservative substitutions, including R>K vs. K>R, G>A vs. A>G, Y>F vs. F>Y, and T>S vs. S>T, showing that substitution classes are not simple reciprocal exchanges. Several observed classes have human-relevant precedent. The frequent A>S class is consistent with the conserved challenge of preventing serine-for-alanine mistranslation by alanyl-tRNA synthetase editing^78^, whereas alanine-producing substitutions such as G>A, E>A, V>A, and T>A are supported by natural human tRNA variants that misincorporate alanine at glycine, glutamate, valine, or threonine codons ^79,80^. Less frequent classes in our data also map onto known human mistranslation channels, including F>S from an endogenous human tRNA-Ser variant^81^. At the aggregate level, the most common origin residues were A, V, L, T, E, S, and G, whereas the most common mistranslation destinations were N, I, S, V, T, G, D, and A. Origin totals combine opportunity in the detectable proteome with error propensity, while destination totals highlight recurrent mistranslation products (Supplementary Table 6).

To probe proteome plasticity and assess the functional relevance of non-genetic substitutions in the biological context we investigated four MS datasets of human cancer-derived cells cultured under tryptophan, tyrosine, leucine, or arginine depletion ^9,50,51^. Using our substitution detection workflow, we identified both previously reported and novel substitutions under these conditions. We confirmed reported increase in W>F and Y>F under tryptophan and tyrosine depletion, respectively (Fig. 4c, left) ^9^. We also corroborated the reported increase in R>K, R>C, R>H, and R>S substitutions under arginine depletion, extending the original analysis, previously focused on arginine variation in a single protein, to a proteome-wide scale (Fig. 4c, top right). Notably, leucine depletion led more frequently to misincorporation of phenylalanine or valine at leucine positions (L>F and L>V; Fig. 4c, bottom right) in comparison to negative control. Additionally, leucine starvation also induced a prominent M>T substitution that was not detected in any non-starved control replicates (Fig. 4c, bottom right). Although M>T has been flagged as a potential sample preparation artifact associated with iodoacetamide (IAA) alkylation, its marked and specific enrichment in all replicates of leucine-starved cells is more consistent with a biological origin ^87^. Finally, W>D was detected across all amino acid depleted and non-depleted conditions, but a selective increase in W>D frequency was observed only under tryptophan/tyrosine starvation (Fig. 4c, left), suggesting that W>D may play a distinct physiological role under specific stress conditions. All detected substitutions in starvation datasets are reported in Supplementary Table 7 and complete 20×19 confusion matrices in Extended Data Fig. 5 (R starvation), 6 (L starvation) and 7 (W/Y starvation).

To rule out a chemical or sample-handling origin for the W>D substitution, we performed tryptic digestion of HeLa cells in H₂¹⁸O. Of 38 W>D events identified in the labelled digest, none carried an ¹⁸O incorporation, indicating that these substitutions did not arise during sample preparation. Independent support was also obtained from cross-dataset comparison: 14 of the 38 W>D bearing peptides were also identified in healthy tissues (Fisher exact test, p = 3.1×10⁻¹⁰) and 8 in tryptophan/tyrosine starvation datasets (Fisher exact test, p = 4.8×10⁻⁷), an overlap inconsistent with stochastic oxidation and consistent with *bona fide* biological substitution (Supplementary Table 8, overlaps/detections; Extended Data Fig. 8, spectra).

We applied an analogous strategy to the C>A substitution, using the deuterated-water protocol described by Wang and colleagues on labelled HeLa lysate ^82^. In parallel, we analysed HEK293 and E. coli proteomes to test the kingdom specificity of C>A reported by Sun and colleagues^88^. No C>A events were detected in any of the three samples. We therefore conclude that the vast majority of C>A events in our healthy-tissue dataset arise from TCEP-induced desulfurization during sample preparation rather than from biological mistranslation. Next, we investigated the biological and evolutionary distances between the codons of the reference and substituted residues in substitution cases lacking genetic support. We examined whether the substitution patterns in non-genetic variants are distinct from those originating from mutations. To quantify these differences, we calculated the minimal nucleotide distance between the codons of the reference amino acid and the hypothetical coding sequence for the observed alternative amino acid. For instance, for a putative substitution from glycine (G) to serine (S), the distance is 1 because among the six serine codons, two (AGU and AGC) differ by only a single nucleotide from a corresponding codon for G. We will refer to substitutions within mutational distance 1 as near-cognates and the rest as non-cognates. We found that, among substitutions with RNA-seq support, the minimal nucleotide distance is 1 in most cases (5,001out of 5,130). By contrast, variants without RNA support frequently showed more distant substitutions - 2 nucleotides in 30% cases (1,578 out of 5,216) and occasionally 3 (119 out of 5,216, ∼2%) nucleotides from the reference codon (Fig. 4d). Near-cognate non-genetic substitutions appear to be more abundant in comparison to non-cognate substitutions reflecting higher disruptive potential of the distant amino acid substitutions on protein structure and stability (Fig. 4d)^7^. Consistent with these observations, the distribution of BLOSUM62 scores^89^ (a score representing evolutionary acceptance rate of the amino acid substitution between homologous protein segments) for substitutions lacking genetic support was shifted toward more negative values (i.e. lower observed rate of evolutionary substitutions), indicating a larger functional distance between the reference and substituted residues (Fig. 4e).

Finally, we assessed the substitutions without RNA-Seq support for their potential occurrence in the human population as per gnomAD 4.1 statistics. We evaluated substitutions with a sequencing coverage of 10 reads or more, all supporting the reference variant but not the alternative. Similarly to our previous analyses on genetically encoded substitutions, we observed a significant correlation (Pearson 0.55 p=1.88×10^-20^) between substitution abundance ratio and its allele frequency in human population (Fig. 3c,f).

### Abundant non-genetic substitutions provide means for functional protein diversification

We analyzed the potential functional consequences of mistranslations by means of AlphaMissense pathogenicity score (Supplementary Table 9)^90^. High AlphaMissense pathogenicity score of the substitution indicates a disturbance of a functionally important site with potential functional outcomes such as change of protein structure, modulation of the protein activity or gain of a new function. Considering that alternative protein variants mostly represent the minority of the gene’s total protein product, we interpret their pathogenicity scores as a potential modulation of the main protein function, implemented by the reference proteoform. The higher pathogenicity score serves as a proxy for a potential “disruptiveness” of a given substitution. We performed a separate functional analysis on non-genetic and genetic substitutions in our dataset. We specified a minimum RNA sequencing coverage threshold per substituted position as 10 reads. We observed a negative correlation between predicted pathogenicity and the substitution abundance ratio in both genetic and non-genetic variant groups, indicating that more disruptive substitutions tend to be present at lower levels in the proteome (Fig. 5a). For genetic variants, this relationship likely reflects the greater disruptive potential of somatic substitutions (typically characterized by lower abundance ratios) relative to germline variants. The analogous trend among non-genetic variants is consistent with either (i) preferential synthesis of non-pathogenic variants or, more plausibly, (ii) enhanced degradation of more deleterious protein variants. Notably, correlation with predicted pathogenicity is weak and represent only one of multiple determinants of substitution abundance. Moreover, non-genetic substitutions classified as pathogenic may not necessarily impose a comparable fitness burden, as they are incorporated into only a fraction of a protein’s molecules rather than uniformly affecting all copies, as implicitly assumed by AlphaMissense. Such partial incorporation could, in principle, expand functional diversity while avoiding the full fitness cost of a fixed genetic mutation.

**Figure 5.**
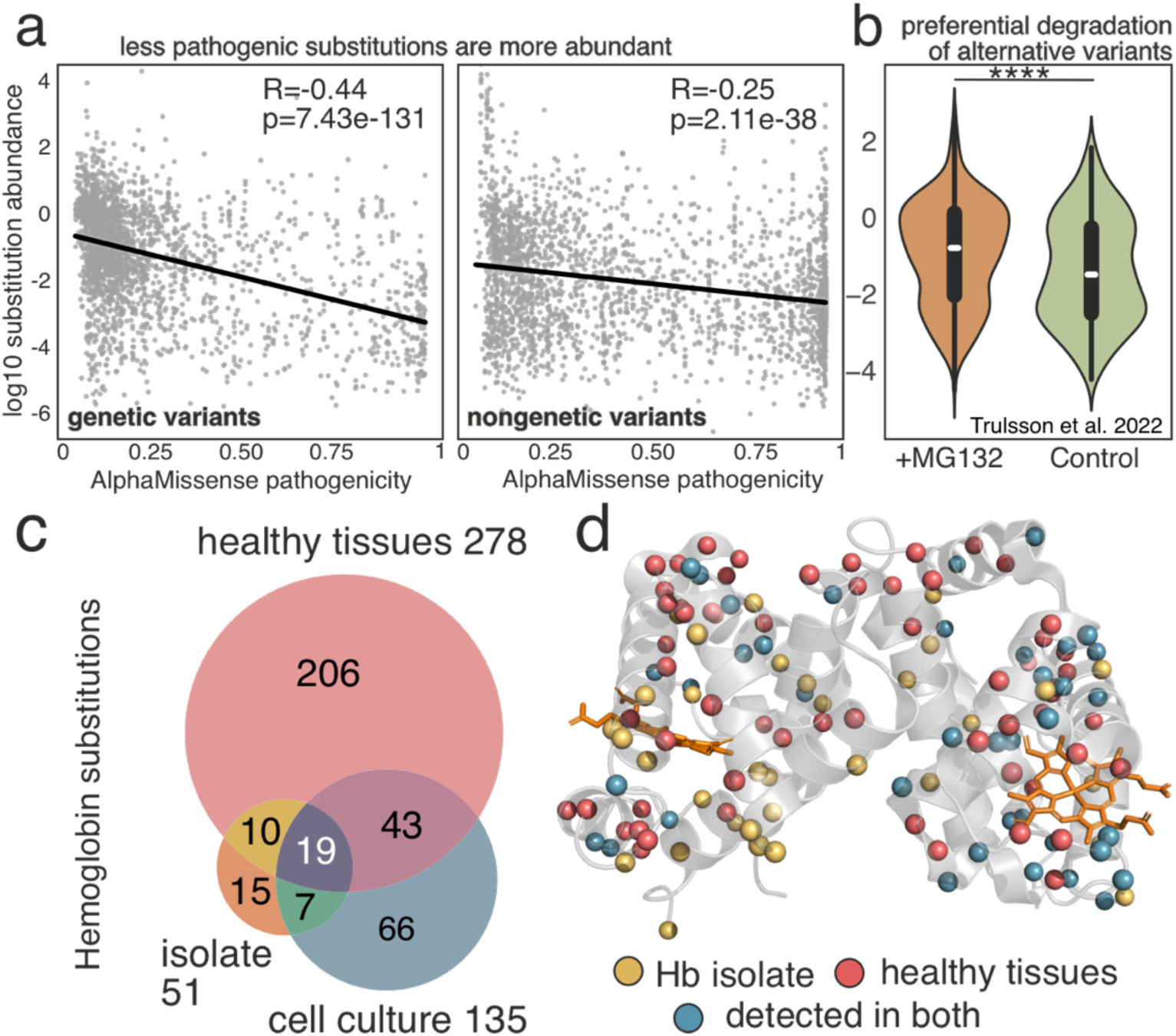
Functional characteristics of amino acid substitutions. (a) AlphaMissense pathogenicity scores plotted against substitution abundances of genetic (left) and non-genetic (right) variants. (b) Distributions of alternative variant substitution abundances in proteasome-inhibited (left) and non-inhibited conditions (right). (c) Venn diagram of hemoglobin substitutions identified in three datasets. Red – substitutions found in healthy tissue dataset, orange – in purified hemoglobin isolate, blue – in in vitro differentiated hematopoietic precursor cell culture. (d) Positions with identified substitutions in human hemoglobin. Red spheres – variants found only in healthy tissue dataset, yellow spheres – variants only detected in purified hemoglobin, blue spheres – substitutions identified both in healthy tissues and purified hemoglobin.

To examine whether alternative variants are subject to selective degradation, we leveraged a dataset from Trulsson et al. (2022) in which osteosarcoma-derived U2OS cells were cultured in the presence or absence of the proteasome inhibitor MG132^52^. If alternative proteins were as stable as their reference forms, their relative abundance would be largely unchanged upon proteasome inhibition. Instead, we observed a global increase in the abundance of alternative protein variants following proteasome inhibition, indicating preferential proteasome-dependent turnover (Fig. 5b, Extended Data Fig. 9 for replicate-separated abundance distributions). This suggests that the steady-state levels of alternative proteins may be substantially higher in the absence of active proteasomal degradation. All substitutions detected in the Trulsson et *al.* dataset and their corresponding abundance ratios are listed in Supplementary Table 10.

Further focusing on functional consequences of the amino acid substitutions, we searched the expanded dataset of substitutions without RNA-seq support for variations within the functional sites of proteins. We identified 85 unique positions in 22 proteins with substitutions within the annotated functional sites in UniProt database (Supplementary Table 11, only keywords ”site”, “binding site” and “active site” were considered, sample spectra Supplementary Fig. 1). We hypothesized that substitutions of residues in the vicinity of annotated functional sites would also modulate protein activity. To assess whether substitutions preferentially occur near functionally important residues, we performed a randomization test. For each observed substitution, we computed its linear distance (number of residues between the sites along the protein sequence) to the nearest annotated functional site and summarized these distances by the median across substitutions. We then compared the observed median distances to a null distribution generated from 20,000 simulated datasets in which substitution positions were randomized within the MS-observed regions of the corresponding protein sequences. Substitutions in our dataset were significantly more clustered near annotated active sites than expected by chance (p = 5×10^-5^), whereas no significant clustering was detected around annotated binding sites (p = 0.06). This enrichment near active sites is consistent with the possibility that some mistranslations are functionally consequential, as substitutions close to catalytic residues are more likely to affect enzymatic activity. These results therefore support the idea that mistranslation can broaden the functional repertoire of the proteome by generating low-abundance variants with altered biochemical properties. Notably, many substituted functional-site residues involved C, H, M, and W - substitutions we previously flagged as “potentially non-biological.” These substitution types were excluded from the randomization analysis. Nevertheless, because we cannot rule out a biological contribution, we report them alongside other functionally relevant substitutions in Supplementary Table 11, with uncertain origin indicated in the “ambiguous” column.

Next, we examined whether some of the amino acid substitutions without RNA-seq support (disregarding their functional annotation) in the unfiltered dataset, appear regularly across different tissues and individuals. Recurrent appearance of the substitution across samples (*i.e.* across tissues and individuals) suggests its potential functional significance. Further, such recurrence bolsters our confidence for its non-genetic origin, as the probability of absent sequencing evidence for a recurrent mutation in unrelated individuals is restrictively low. We gathered a set of 163 proteins and 392 unique substitutions which are repeatedly detected in 5 or more different tissues from different individuals (Supplementary Table 12, sample spectra in Supplementary Fig. 2-6).

As expected, a larger number of distinct substitutions was observed in highly abundant proteins, where detection of low-frequency alternative forms is technically more tractable. For example, we detected 227 and 44 unique variants in hemoglobin and GAPDH, respectively; of these, 46 hemoglobin substitutions and 10 GAPDH substitutions were observed recurrently in five or more individuals (Supplementary Table 12). To assess whether the increased number of variants in abundant proteins could reflect ambiguous spectral assignments in complex whole-tissue matrices, we expressed these proteins in a purified form and subject them to a new MS measurement and analysis. We searched for alternative protein variants in these enriched or purified protein preparations. Specifically, we analyzed (i) 121 immunoprecipitated proteins from HEK293 cells generated as part of the BioPlex project and (ii) four highly purified proteins (hemoglobin, GAPDH, PRDX6, PARK7, >98% purity) produced recombinantly in, or isolated from, human cells ^91^. To test whether mistranslation is detectable on a sequence template that is orthogonal to the endogenous human proteome, we analyzed a recombinant antibody expressed and purified from HEK293 cells whose amino acid sequence shares no substantial similarity with any protein in the human reference proteome (Supplementary Fig. 11,13,15). In comparison to healthy tissues samples, all these experimental datasets employed distinct sample preparation protocols and relatively fresh material, minimizing the likelihood of artifactual substitutions.

Across the 121 BioPlex immunoprecipitations, we detected 2,748 substitutions (median 21 per protein), including 54 variants that were also observed in our healthy-tissue analysis. In our analysis of highly purified proteins, we detected 60, 31, 27, and 27 substitutions in hemoglobin, GAPDH, PARK7, and PRDX6, respectively; of these, 37, 5, 3, and 4 were also detected in healthy tissues. All the overlaps were modest but statistically significant as per Fishers exact test (p-values 4.12×10^-21^/1.24×10^-9^ for hemoglobin subunits, 0.02 for PARK7, 3.2×10^-5^ GAPDH and 4.2×10^-5^ PRDX6). Notably, both the BioPlex samples and our purified-protein preparations were freshly generated and MS sample preparation did not employ desulfurization-promoting protocols. Nevertheless, we again observed abundant C>A, C>S, W>D, H>D, and M>D substitutions, supporting the possibility that at least a subset of these events may arise from *bona fide* biological processes. All substitutions identified in enriched samples from BioPlex project and purified proteins are reported in Supplementary Table 13 and 14 respectively and sample mirror spectra of co-identified peptides in healthy tissue samples and purified proteins on Supplementary Fig. 9 and 10. The limited overlap of substitution identifications across samples likely reflects current methodological constraints rather than biological variability. We expect the true extent of the substitution landscape to be revealed more comprehensively as instrumentation sensitivity and computational tools for open/exploratory searches continue to improve.

Further, mass spectrometric analysis of the recombinant antibody recovered 45 substitutions that could not be explained by either the construct’s coding sequence or contamination from endogenous human or bovine immunoglobulins (IgBLAST, Supplementary Fig. 11,13). This orthogonal experiment, performed in a non-endogenous background, confirms that mistranslation by the human ribosome is persistent and includes non-trivial substitution classes with mutational distance >1 supported by high-quality MS/MS evidence (Extended Data Fig. 10, Supplementary Fig. 12,14 and Supplementary Table 14).In tissue proteomics, including the analyses reported here, hemoglobin is typically among the most abundant proteins due to tissue vascularization and residual blood content. As noted above, we detected 227 unique substitutions across the α- and β-subunits of hemoglobin (Fig. 5c,d). Some of these distinct substitutions were detected in multiple tissues yielding total 530 hemoglobin substitution events in the whole dataset (Supplementary Table 1). Hemoglobin function depends on an intricate allosteric network that enables the heterotetramer to transition between low- and high-affinity oxygen-binding conformations. Consequently, substitution in even a single subunit could yield hybrid tetramers with altered biological and physicochemical properties ^92^.

Among 227 unique, technically non-ambiguous hemoglobin substitutions, 27 matched exactly to previously characterized clinically relevant hemoglobin variants, including 20 associated with increased and 7 with decreased oxygen affinity (Supplementary Table 15). For these variants, the functional effects are supported by prior experimental characterization of the same amino-acid substitutions. A further 73 variants affected residues that are clinically relevant in hemoglobin but involved different amino-acid exchanges. We describe these as position-level functional candidates rather than validated functional variants. Their prioritization is based on same-residue clinical relevance and biochemical severity features (e.g. volume change upon substitution, change in amino acid class upon substitution etc.), but their precise functional effects remain inferential (Supplementary Table 16).

We hypothesized that some of this diversity could arise genetically from mutations accumulating across the limited number of hematopoietic lineages that ultimately generate mature red blood cells. Consistent with this possibility, among 530 non-unique hemoglobin substitutions detectable by MS, 82 (15%) were observed at low levels in RNA-seq data. However, the remaining MS-detected substitutions lacked corresponding sequencing support. To minimize potential confounding effects from genetic heterogeneity of the analyzed material and from red blood cell aging, we additionally analyzed a dataset of *in vitro*–differentiated erythroid precursors from Karayel et al. (2020)^93^. In this independent dataset, we detected 135 unique hemoglobin substitutions, 62 of which overlapped with those observed in healthy tissues (Figure 5d, Supplementary Table 17), representing a significant overlap (hypergeometric p-value = 4.06×10^−46^) (Supplementary Fig. 7; spectra). These results independently confirm that hemoglobin exhibits extensive substitution diversity *in vivo*, although the underlying mechanisms driving this diversification remain unclear.

In all these analyses we used the MS-supported substitutions without the RNA-seq evidence as a proxy to derive the general characteristics of what are hypothesized to be mistranslations. We are aware that due to the RNA-seq coverage limitations we cannot completely exclude the cases of somatic mutations within the tissue samples. However, we assume that the averaged trends over different amino acid substitution patterns, their pathogenicity and allele frequencies would largely reflect the tendencies of protein mistranslation as it represents a major source of protein heterogeneity in cells^7,37^. Additionally, the existence of recurring substitutions in samples from non-related individuals allowed us to rationalize on distinct roles of non-genetic substitutions in human.

## Conclusion

In conclusion, our data reveal that unity and multiplicity coexist within the proteome: the genome’s unique signature is represented not just by a single encoded form, but rather by a spectrum of functionally relevant proteoforms that blur the line between individual and population. Our study, therefore, provides a unique perspective on an additional layer of genetic design complexity, highlighting the existence of a functional, recurrent, and evolvable protein quasi-species within the human proteome^94^.

## Methods

### RNA-seq analysis

Raw RNA sequencing reads in fastq format were downloaded from Array Express (http://www.ebi.ac.uk/arrayexpress/experiments/E-MTAB-2836/) and their correspondence with the MS data was verified through Table EV1, tab “Sample Information” in Wang et *al*. publication^49^. Fastq files were aligned against ensembl CDS reference sequences (r112, https://ftp.ensembl.org/pub/release-112/fasta/homo_sapiens/cds/) using BWA^95^. Resulting files were converted, sorted and indexed using SAMtools^96^. Unmapped reads as well as supplementary, secondary and duplicated alignments were discarded. Observed codons at the genomic locations of each amino acid substitution were counted for each sample separately, and counts were summed per tissue. Alignments that contained N at the substitution position or an indel prior to it were removed.

We considered a substitution confirmed on a genetic level if observed number of reads with mutation could not arise from the DNA polymerase used in the sequencing library preparation. Specifically, we compute a one-sided binomial tail probability:

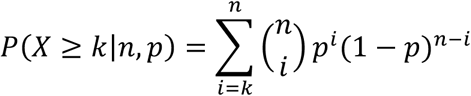

k = number of reads reporting the specific non-reference base that would produce the amino-acid substitution,
n = total RNA-seq read depth at that genomic position,
p = per-read probability of observing that specific base due to error. We define p as:

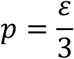

ε is the assumed per-base error rate (PCR/sequencing, 0.001 used in this manuscript based on reported values from ^97^ ), and division by 3 reflects that (under a simple symmetric model) an error at a base can convert to any of the other three nucleotides, but we are testing one specific alternative nucleotide.

Finally, we apply Benjamini–Hochberg correction across all substitutions tested to control FDR. The binomial test is performed at the RNA-seq level, the proteomics observation is not incorporated into the p-value calculation to avoid inflation of the dataset with the false positive genetic identifications.

### MSFragger search for the substituted candidates

Raw data for the analysis were obtained from Proteomic identification database (PRIDE ID PXD010154 for healthy tissue data, PXD027328 for degradomics dataset, PXD017276 for *in vitro* differentiated RBC precursors, PXD043612 for arginine starvation dataset, PXD044085 for leucine starvation dataset, PXD028921 for tryptophan/phenylalanine starvation dataset, ), BioPlex APMS datasets from MassIVE MSV000080679 repository and immunoglobulin profiling data from Zenodo database reference 10.5281/zenodo.14835053. and processed by FragPipe 22.1.07^61^. To separate search spaces for the reference and substituted peptide searches, a two-stage FDR-control protocol described previously was implemented. ^57^. Raw data were searched using reference proteome database. Human protein sequences were taken from the Homo sapiens GRCh38 Ensembl release 112 peptide FASTA, together with the matched CDS/transcript FASTA, because our pipeline required codon-level mapping of candidate mistranslated residues. First search was performed using default parameters with MSBooster and IonQuant quantification activated with default settings^55,61^. Methionine oxidation was set as variable and cysteine carbamidomethylation as fixed modification. In case of degradomics datasets from Trulsson et *al.* additional variable modification of mass 114.04293 was set on lysine. "Write sub mzML” option in the Run tab of FragPipe was allowed in order to separate unidentified spectra and generate a workflow file for the following second search^57^. Automatic selection of best predictors of retention time and MS2 fragmentation spectra was chosen^98^. IonQuant was used for MS1 quantification with default parameters^62^. Match between runs was not activated.

Identified peptides from the first search were in silico mutagenized using custom python script and used as a reference database for the following search. The workflow file and new raw data in mzML format generated by a first search were used to perform a second search. The parameters were left default except: “Protein Digestion” on the MSFragger tab of FragPipe was changed to “nocleavage” to prevent further cleavage of the searched peptides, N-terminal methionine clipping was deactivated and minimum number of scans and isotopes for IonQuant quantification was decreased to 1. MSBooster was activated and retention time and MS2 fragmentation spectra predictors were set to the predictors selected in the first search (DIA-NN for the retention time prediction^99^ and AlphaPept MS2 Generic for the fragmentation spectra prediction^100^). In case of ^18^O labeled experiment, peptides were searched with allowed variable +2.0043 or +4.0085 Da modification on C-teminus and +4.0085 Da variable modification aspartate. In case of D_2_O labeled experiments, peptides were searched with allowed variable +1.0073 or +2.014 Da modification on C-/N-termini and +1.0073 Da variable modification on alanine.

### Validation of peptide candidates

First, spectral index was created from the MSFragger first search output - unidentified raw spectra in mzML format. Index was created as per PepQuery documentation. PepQuery was run in PSM validation mode in parallel for all identified PSMs on computational cluster. Input files for each parallel job consisted of tab separated values of the evaluated peptide sequence and its spectrum title reported in the psm.tsv file from the MSFragger first search. The validation search was set as follows: 20 ppm accuracy (-tol 20), 0-3 isotopic errors (-ti 0,1,2,3), fragment ion m/z tolerance of 0.05 Da (-itol 0.05) for Orbitrap and 0.6 for ion trap data. Enzyme cleavage was set to Trypsin (no P-rule, -e 2) and maximum two missed cleavages were allowed (-c 2). Search was not performed in the fast mode and scoring function was set to Hyperscore (-m 1). Validated peptide was required to have higher score than any of the competitors (-hc). Validation included competition with all possible post-translational modifications from UniMod database^101^ as well as with single amino acid substitutions of the reference proteome peptides (-aa). The latter option caused that majority of the validated peptides were not reported (as they indeed represented a substituted reference peptides which are competed against in PepQuery validation) and were recorded in the ptm_detail.txt file together with all other potential PTM’s and reference peptide variants. All reported variants for the tested peptides were extracted from the ptm_detail.txt. Peptide was considered validated if it had the highest score in comparison to the other candidates. PSM’s with equally scoring candidates were discarded. After the validation, all spectra were annotated graphically by PDV version 2.1.2 and b/y ion series were output in textual format. Only PSM’s with at least two MS2 fragments per validated substituted position were considered for the further analysis. Additionally, substitutions found on the N-terminus of the peptide were not retained due to the low confidence of b1 ion identification^65^. Next, each spectrum of the identified alternative peptides was matched with its reference proteome counterpart using PepQuery without considering precursor mass (i.e. only on MS2 level). Spectra matching the reference peptides with higher score than alternative peptide matches were removed. Finally, all PSM’s were processed by pyAscore algorithm with precision setting 0.01 Da for orbitrap and 0.6 Da for ion trap data. Only hits containing confident localization on the assumed substituted position were retained.

### Processing and filtering of the validated substituted peptides

For quantification, peptides were further filtered with the custom python script. Substitutions located on any position within 4 residues from the N’ or C’ termini were not considered due to potential trypsin cleavage bias^66^. Only correctly and singly mis-cleaved tryptic peptides were retained. Substitutions originating from or resulting in lysine or arginine were not considered due to quantification bias from the removal of existing or creation of a novel trypsin cleavage sites. For substitution abundance calculation, maximum observed intensity of the reference and alternative peptides of the same charge state was utilized. If reference and alternative peptide were found in several charged states, abundance ratio was calculated for every charged state separately and their median was taken to yield the final substitution ratio. This set of filtered substitutions was called ”filtered data” and used for all analyses involving substitution abundance ratio.

### MaxQuant dependent peptide search and validation

Raw data were downloaded from Proteomic identification database (PRIDE ID PXD010154) and searched against Ensembl hg38 v112 proteome database using MaxQuant 2.4.13.0^102^. Dependent peptide search option was activated, match between runs deactivated. Carbamidomethylation of cysteine was set as fixed modification and methionine oxidations as a variable modification. AllPeptides.txt output table was processed by custom python scripts published by Mordret et *al*^7^. Raw files were converted to mzML format using ProteoWizard^103^ and spectrum index for the following PepQuery validation was created according to the PepQuery documentation. Detected peptides from the dependent peptide search pipeline were searched again using PepQuery2 with the same settings as described in the methods section ”Validation of peptide candidates” with exception of validation mode, tissue sample-specificity and considering the alternative competitive peptides. Validation mode was set to simple peptide search (instead specific PSM validation). Peptides were queried along all the raw data disregarding tissue specificity. Identified peptides were not scrutinized against possible match with substituted variants of a different reference proteome peptide.

### AlphaMissense analysis

AlphaMissense predictions were obtained at https://zenodo.org/records/8208688 as AlphaMissense_aa_substitutions.tsv file. ^49^Correlation between substitution abundance and their corresponding AlphaMissense pathogenicity was analyzed separately for genetic and non-genetic sets of substitutions.

### GnomAD analysis

Each MS-observed amino acid was assigned a reference genome codon extracted from the CDS file associate with v112 ensembl genome release. Genomic coordinates for each codon corresponding to the MS-observed substitution were extracted with ensembldb utility in R^104^. The table containing genomic coordinates for the MS-observed substitutions was joined with GnomAD v 4.1 exome data downloaded at https://gnomad.broadinstitute.org/dataand containing collection of SNPs, their genomic location and their corresponding population allele frequencies^71^. This joined table was filtered only for such SNPs which would result in MS-observed missense substitution using custom python script. The substitution abundances were evaluated considering their hypothetical allele frequencies within the human population. Three groups of substitutions were evaluated – heterozygotic alleles, somatic variants and variants without RNA support. All variants with sequencing support were filtered towards elimination of substitutions derived by Illumina error. Heterozygotic variants were required that their coding transcripts coverage ratio lie between log10 values between –0.3 and 0.3 corresponding to two-fold difference in mRNA expression and additionally that sequencing coverage of was at least 10. Somatic variants were required to lie between coverage ratio -1.5 and -3.5 corresponding to 3% and 0.03% of substitution-supporting reads within the number of reads covering that position.

### Immunoglobulin analysis

The sequences of proteins with MS-observed substitutions and which associated with GO terms “immunoglobulin complex” and “antigen binding” (GO codes) were analyzed by the IgBlast tool ^105^. Blastp program was used with KABAT as variable domain delineation system ^106^. The tabular output was parsed by custom python script to extract the variable domain regions from the alignment (CDR’s and FR’s). Substitutions with and without genetic support were analyzed for their location within or outside the immunoglobulin annotated region.

### Calculation of amino acid substitution fractions

Amino acid counts from the unique peptides yielded from reference proteome search (first search of the pipeline) for all the analyzed tissues were aggregated together. Next all unique amino acid substitutions originating from each kind of amino acid throughout all the samples were also counted. Substitutions fraction for a given pair of origin and destination amino acid was calculated as a fraction of observed substituted origin amino acids to the total number of observed amino acids of this kind.

### Synthetic peptide analysis

All synthetic peptides were ordered from GenScript (>=98% purity). Their sequences are listed in the Supplementary Table 18. The peptides were dissolved to 1nmol/μL separately and then mixed together to a stock of 10pmol/μL per peptide. The sample was then further diluted to 50fmol/ul, and 9 ul were injected into the instrument. The resulting peptides were analyzed using nanoflow liquid chromatography (nanoAcquity) coupled to high-resolution, high mass accuracy mass spectrometry (Fusion Lumos). Data was searched against the list of peptides using the Byonic search engine and quantified using Skyline.

### Proteomic sample preparation

Pure proteins (GAPDH (MedChemExpress, HY-P78776), PRDX6 (CusaBio, CSB-MP018659HU), PARK7 (HUABio, HUA-HA211158) and Hemoglobin (MP-Bio, MPB-0855914)) were dissolved from lyophilized powder with 5% SDS in 50 mM Tris-HCl. 30ug was used for digestion Samples were prepared according to the previously published protocol optimized for increased digestion efficiency ^107^. Samples were reduced with 5 mM dithiothreitol and alkylated with 10 mM iodoacetamide in the dark. Phosphoric acid was added to the lysates to a final concentration of 1.2%, followed by the addition of 90% methanol in 5 mM ammonium bicarbonate. Each sample was then loaded onto an S-Trap 96-well plate (Protifi, USA). Samples were then digested with trypsin (1:50 trypsin/protein) O/N h at 37 °C. The digested peptides were eluted using 50 mM ammonium bicarbonate; trypsin was added to this fraction and incubated 4hr at 37 °C. Two more elutions were made using 0.2% formic acid and 0.2% formic acid in 50% acetonitrile. The three elutions were pooled and vacuum centrifuged to dry. Samples were kept at −20 °C until analysis. In case of ^18^O and D_2_O labeled experiments, all reagents were freshly dissolved in labeled water.

### High pH Reverse Phase fractionation

UPLC/MS grade solvents were used for all chromatographic steps. 100ug of the whole cell lysates and 27ug of the purified protein digests were fractionated using High Performance Liquid Chromatography (Acquity H Class Bio, Waters, Milford, MA, USA). Mobile phase was: A) 20 mM ammonium formate pH 10.0, B) acetonitrile. Peptides were separated on an XBridge C18 column (3×100mm, Waters) using the following gradient: 3% B for 2 min, linear gradient to 40% B in 50 min, 5 min, to 95% B, maintained at 95% B for 5 min and then back to initial conditions. A total of 90 Fractions were collected (one fraction per 30s) from between 5-50 minutes and combined to a total of 15 fractions. Fractions were then analyzed by MS.

### Mass spectrometry

Samples undergoing DDA acquisition were fractions from High pH RP or the synthetic peptides. UPLC/MS grade solvents were used for all chromatographic steps. Each sample was loaded using split-less nano-Ultra Performance Liquid Chromatography (15 kpsi M-Class nanoAcquity; Waters, Milford, MA, USA). The mobile phase was: A) H_2_O + 0.1% formic acid and B) acetonitrile + 0.1% formic acid. Desalting of the samples was performed online using a reversed-phase Symmetry C18 trapping column (180 µm internal diameter, 20 mm length, 5 µm particle size; Waters). The peptides were then separated using a T3 HSS nano-column (75 µm internal diameter, 250 mm length, 1.8 µm particle size; Waters) at 0.35 µL/min. Peptides were eluted from the column into the mass spectrometer using the following gradient: 4% to 30%B in 55 min, 30% to 90%B in 10 min, maintained at 90% for 7 min and then back to initial conditions. Samples were run in a random order.

The M-Class nanoUPLC was coupled online through a nanoESI emitter (20 μm tip; Fossil Ion Tech, Madrid, Spain) to an Exploris 480 (for 2D fractions) or tribrid Orbitrap Fusion Lumos (for synthetic peptides) mass spectrometer (Thermo Scientific) using a PicoView nanospray apparatus (New Objective).

Data was acquired in data-dependent acquisition (DDA) mode, using a Top-Speed (2sec) method. MS1 resolution was set to 120,000 (at 200m/z), mass range of 375-1500m/z, AGC set to 200% and maximum injection time was set to Auto. Minimum intensity to 5e4, charges limited to 2-6 and dynamic exclusion to 40s. MS2 resolution was set to 15,000, quadrupole isolation to 1.6 m/z, AGC to 75%, maximum injection time to Auto, and HCD to 30 NCE.

## Supporting information

all supplementary tables

## Data availability

Raw data was submitted to PRIDE repository under ID PXD073135.

## Code availability

The substitution search pipeline described here is based on running several third-party freely accessible tools (FragPipe first and second search, PepQuery index building and validation, labile PTM filtering, localization scoring and PDV annotation). We provide a detailed general guidelines on running these software packages, data analysis and evaluation in the methods section. Additionally, we provide example scripts used for complete substitution search and validation on https://github.com/treko90/tarsus.

## Acknowledgements

VT was supported by a postdoctoral fellowship from the Azrieli Foundation. We are grateful to Professor Alexey I. Nesvizhskii, Fengchao Yu and Kevin Yang for providing pre-release version of FragPipe 22.1 with custom modifications and consultations in the initial stages of the project. We are thankful for the Minerva Foundation for establishing the Center for Live Emulation of Evolution in the Lab. YP is an incumbent of the Ben May Professorial Chair. YP is a Kimmel Investigator. We thank Dr. Ziv Shulman and Nachum Nathan for providing the R1 human antibody. We are grateful Hila Levy and Amir Pri-Or from de Botton Institute for Protein Profiling for extensive protein and peptide mass spectrometry work.

## Author contributions

YP/VT designed a study; VT/TL/OA/DM performed the data analysis, VT/YP wrote a draft, OA/DD/DM consulted the study and critically reviewed the text. All authors approved the manuscript.

Supplementary Information is available for this paper.

## Extended data figures

**Extended data Figure 1.**
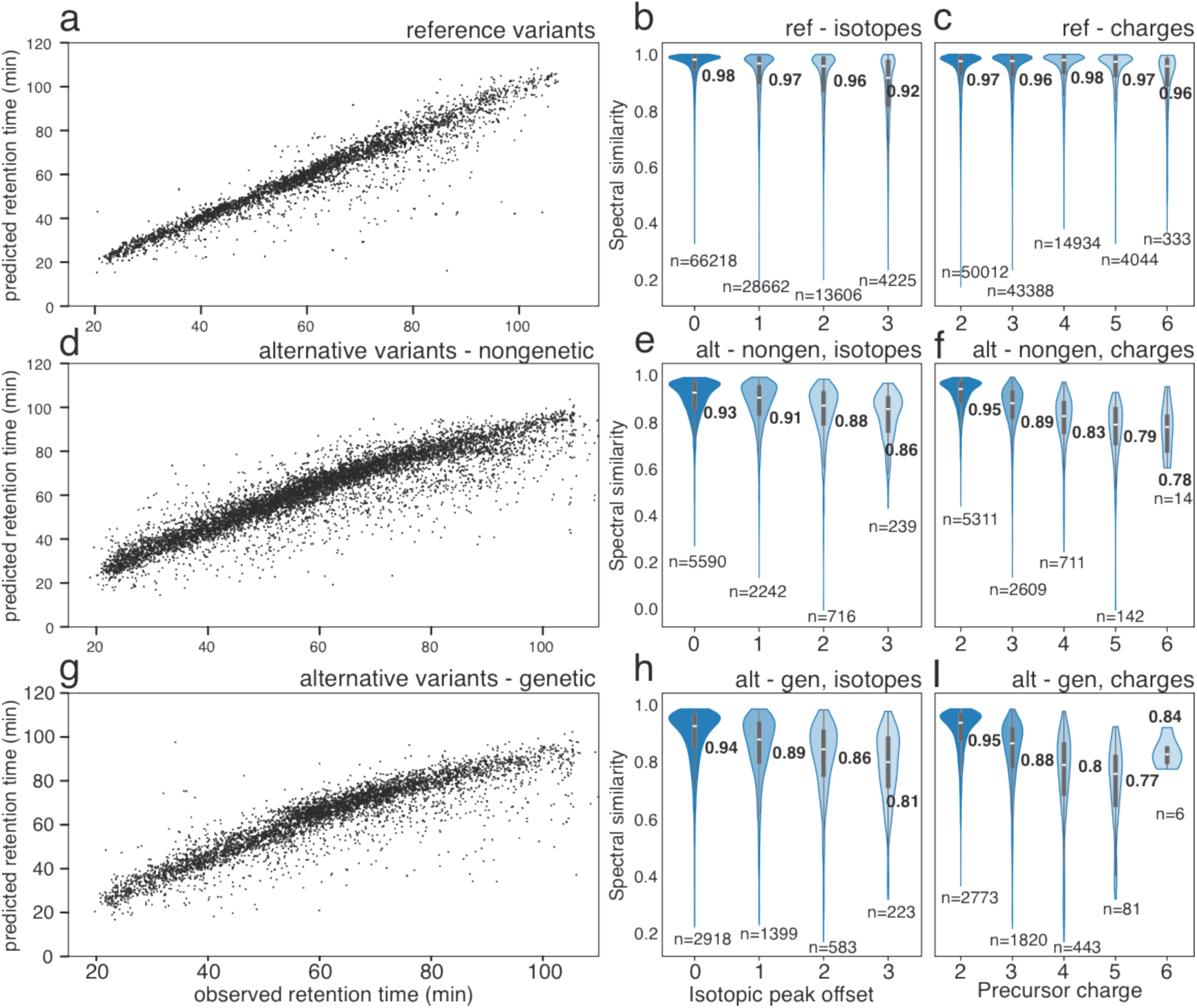
Comparisons of predicted and experimental retention times (left column) and fragmentation spectra separated by distinct isotopic offsets of MS1 precursor (middle column) and precursor charge (right column). Comparisons presented for reference proteome peptides (a,b,c); alternative variants without genetic support (d,e,f) and alternative variants with genetic support (g,h,l)

**Extended data Figure 2.**
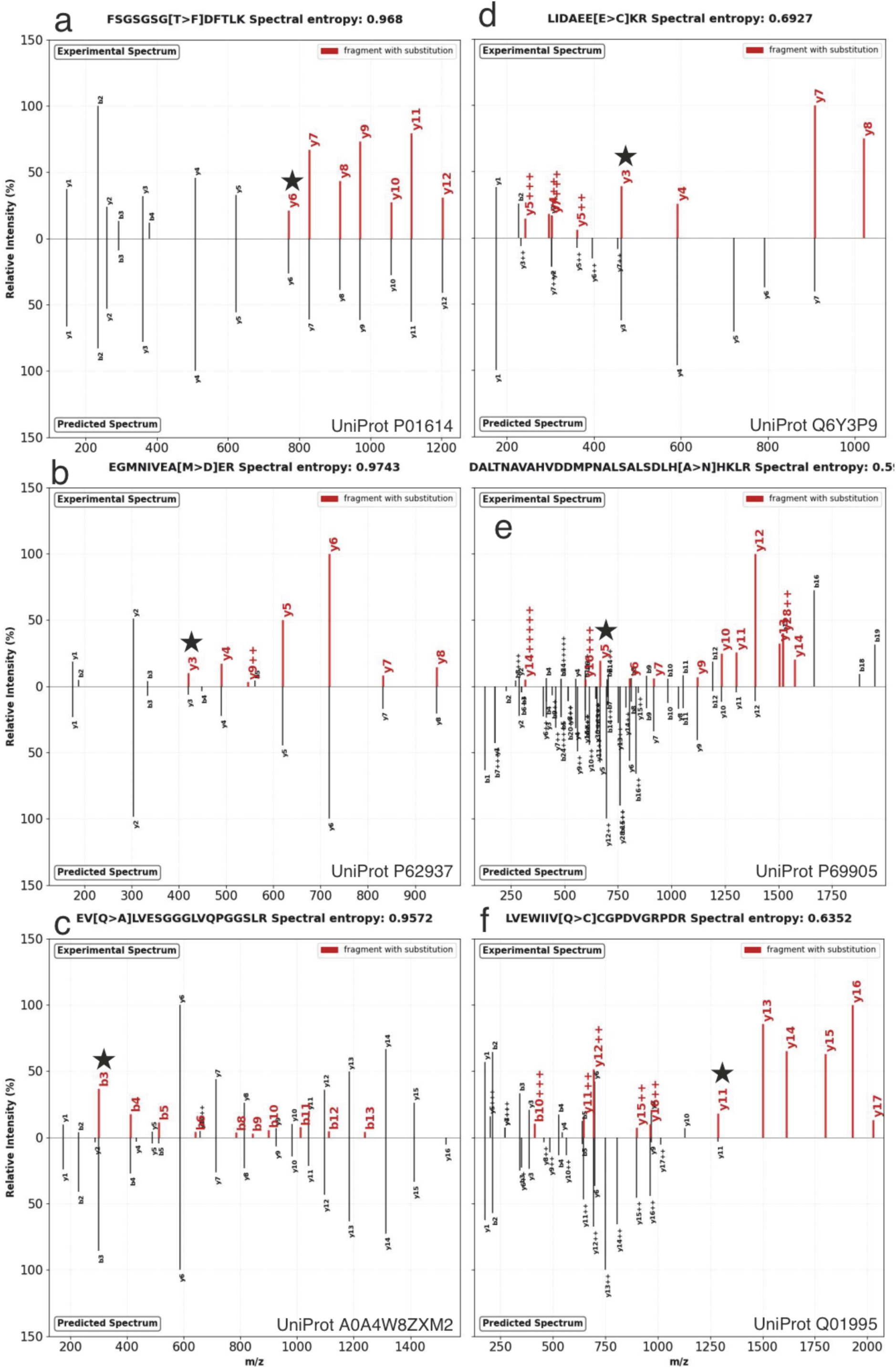
Mirror plots representing predicted (bottom) and observed (top) fragmentation spectra of alternatively substituted precursors. Sample demonstrates spectra of substitutions within mutational distance >1. (a,b,c) Spectra well aligned with the prediction; (d,e,f) spectra with lower similariy with prediction

**Extended Data Fig. 3:**
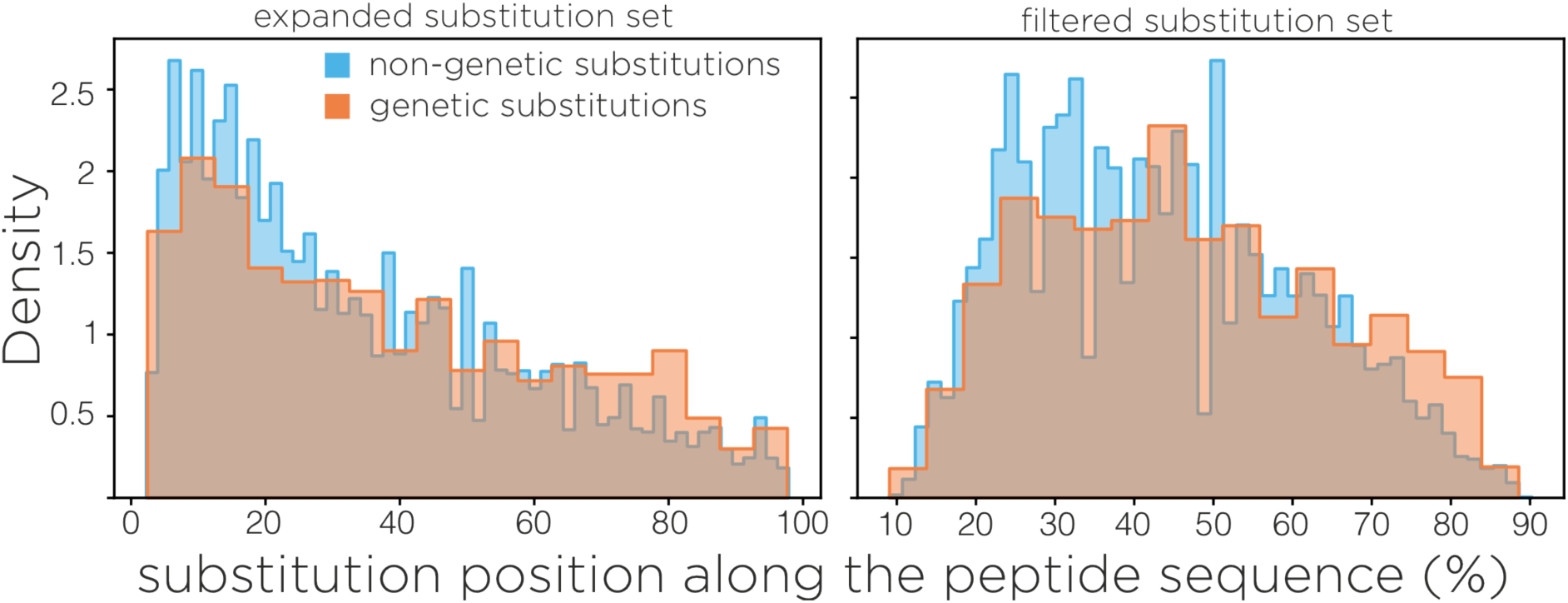
Distribution of positions within the tryptic peptide length where substitution was detected. Right – substitutions within the “extended set”, left – substitutions in “filtered set”. Orange – substitutions mapped to RNA sequencing data, blue – substitutions without RNA sequencing support

**Extended Data Fig. 4:**
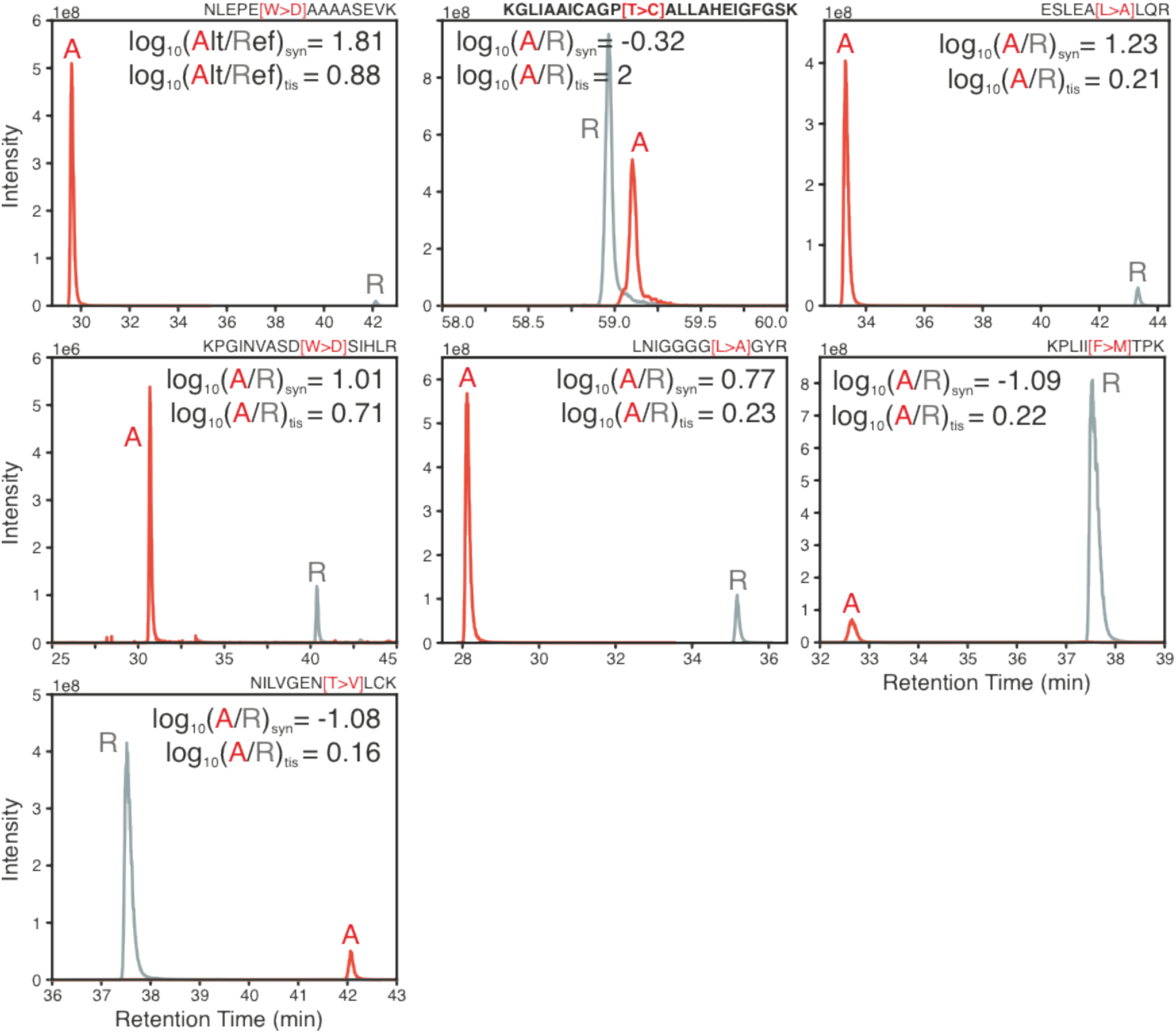
Distribution of positions within the tryptic peptide length where substitution was detected. Right – substitutions within the “extended set”, left – substitutions in “filtered set”. Orange – substitutions mapped to RNA sequencing data, blue – substitutions without RNA sequencing support

**Extended Data Fig. 5:**
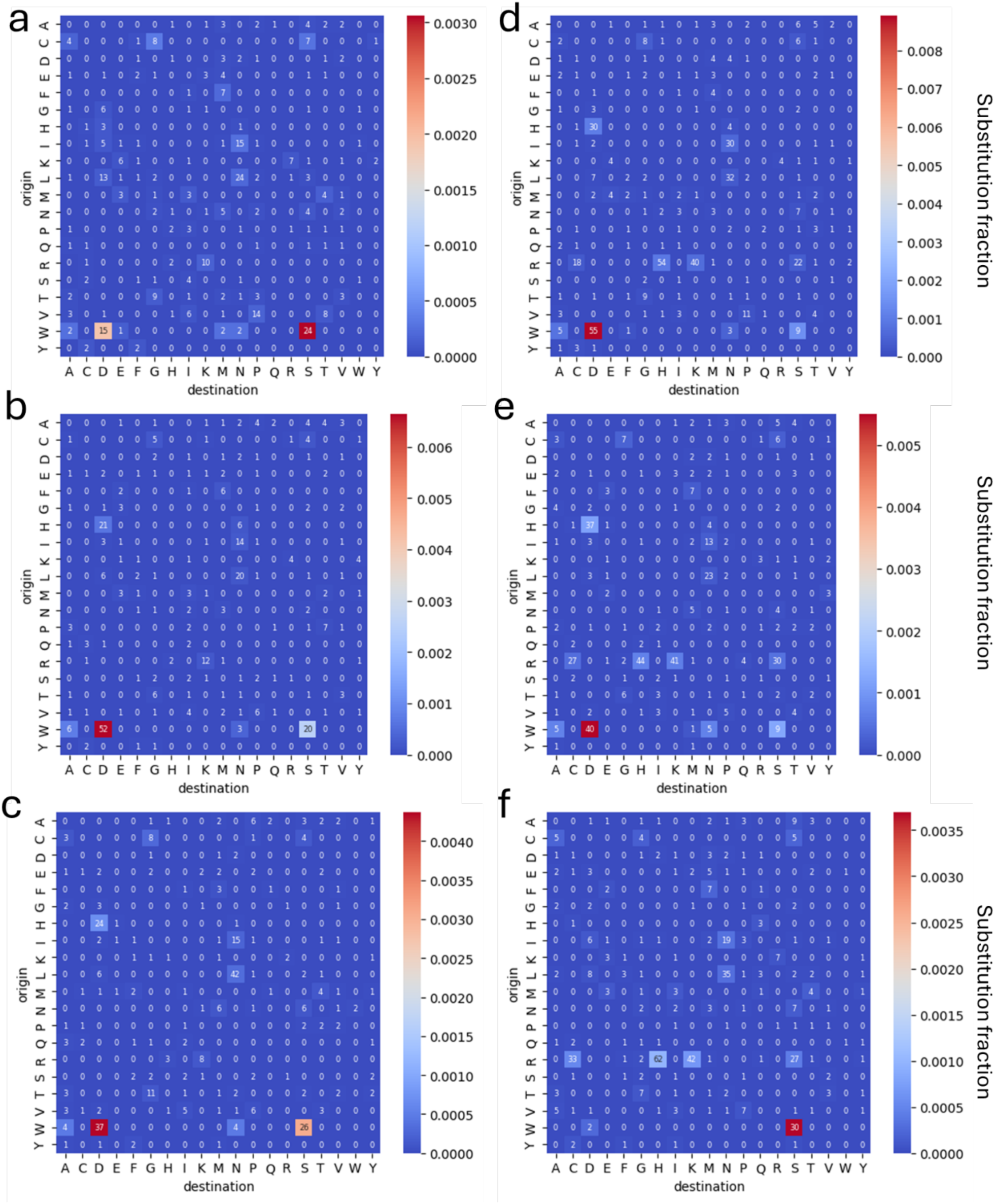
Amino acid substitution matrices with counts (numbers in cells) and substitution fraction (color) of the specific substitutions in control (a,b,c) and arginine depleted (d,e,f) breast cancer MB-231cells. Substitution fraction is defined as ratio of substituted origin amino acids to its total observed count in the MS data.

**Extended Data Fig. 6:**
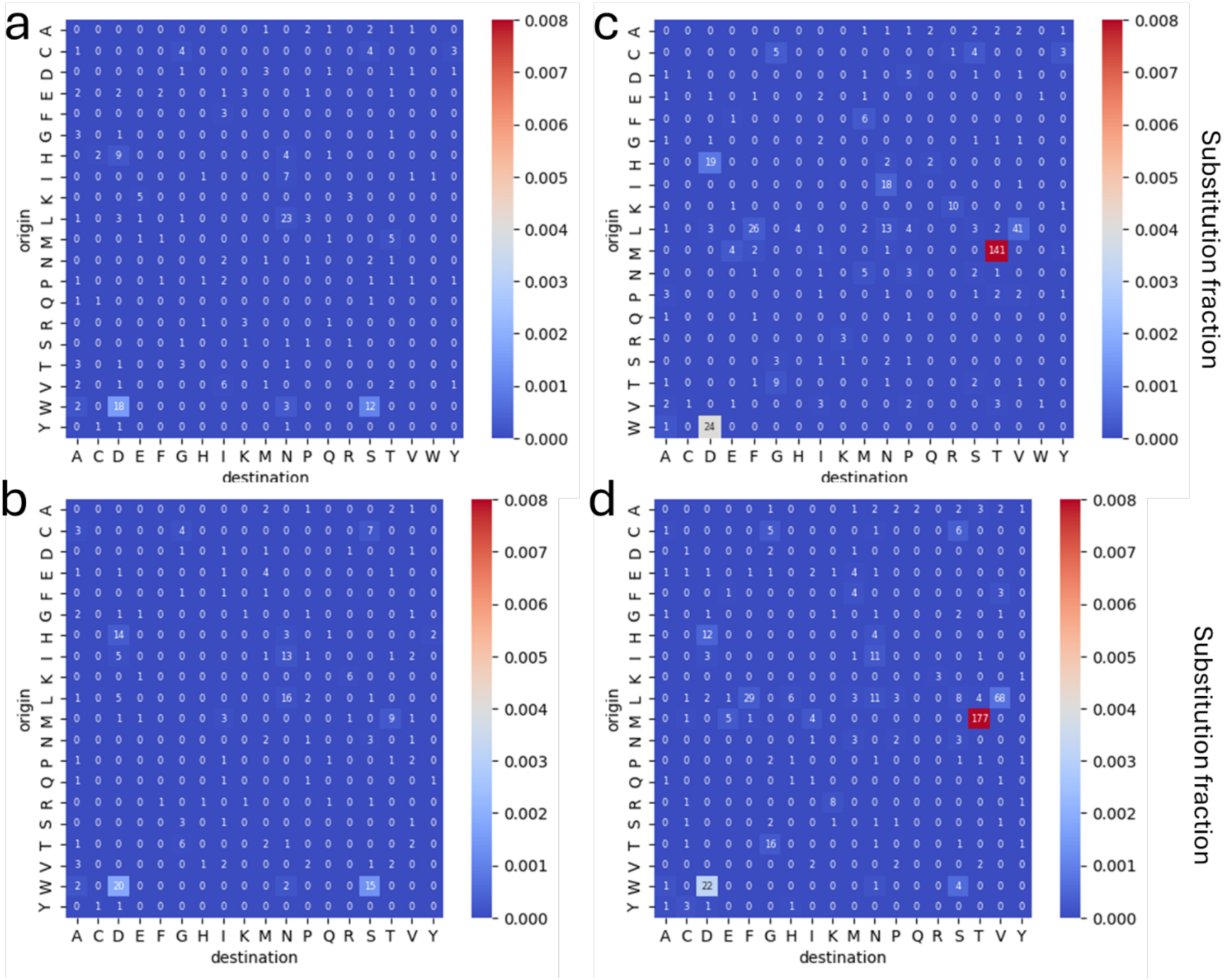
Amino acid substitution matrices with counts (numbers in cells) and substitution fraction (color) of the specific substitutions in control (a,b) and leucine depleted (c,d) breast cancer MB-231cells. Substitution fraction is defined as ratio of substituted origin amino acids to its total observed count in the MS data.

**Extended Data Fig. 7:**
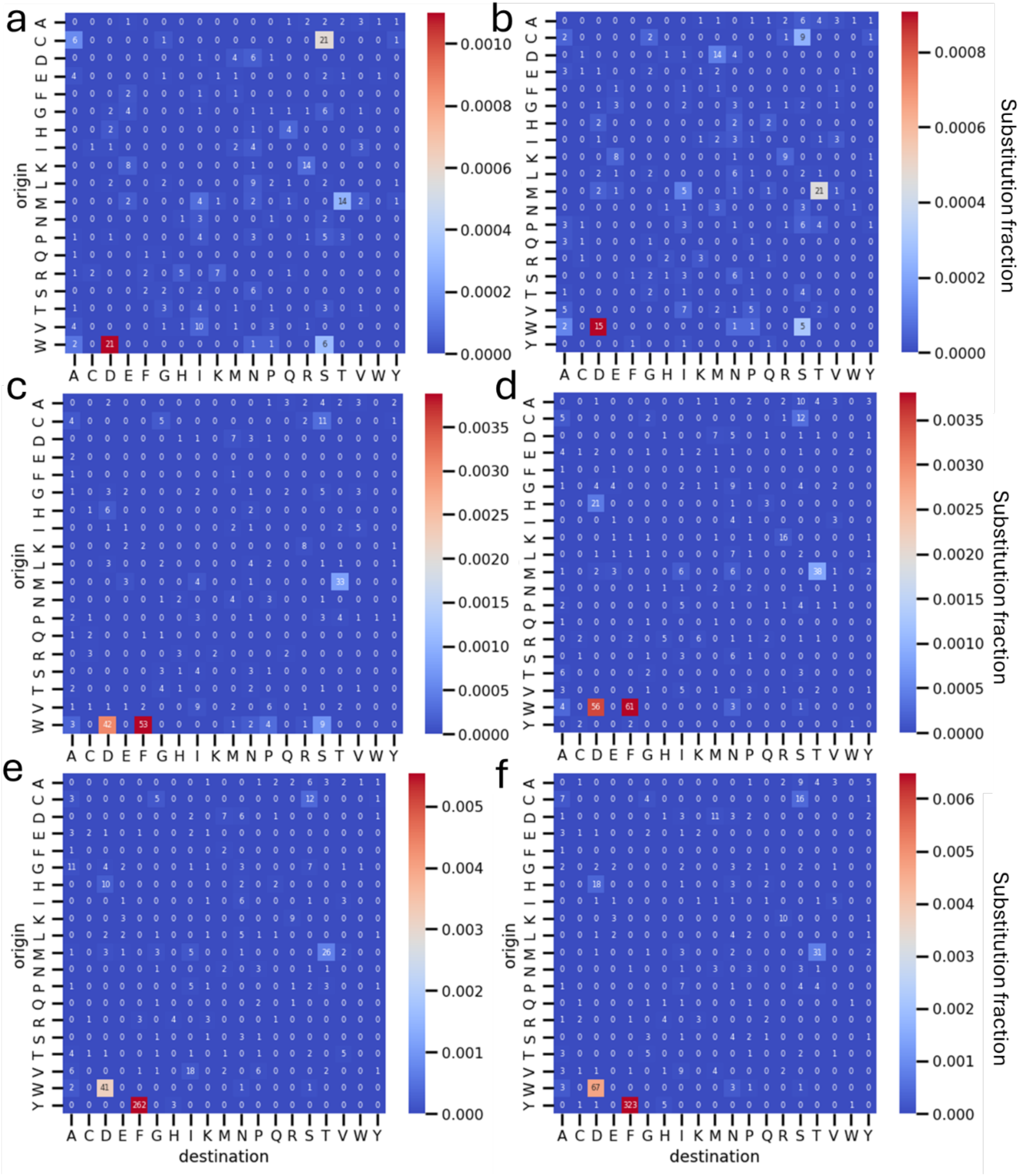
Amino acid substitution matrices with counts (numbers in cells) and substitution fraction (color) of the specific substitutions in control (a,b), tryptophan depleted (c,d) and tyrosine depleted (e,f) melanoma MD55A3 cells. Substitution fraction is defined as ratio of substituted origin amino acids to its total observed count in the MS data.

**Extended Data Fig. 8:**
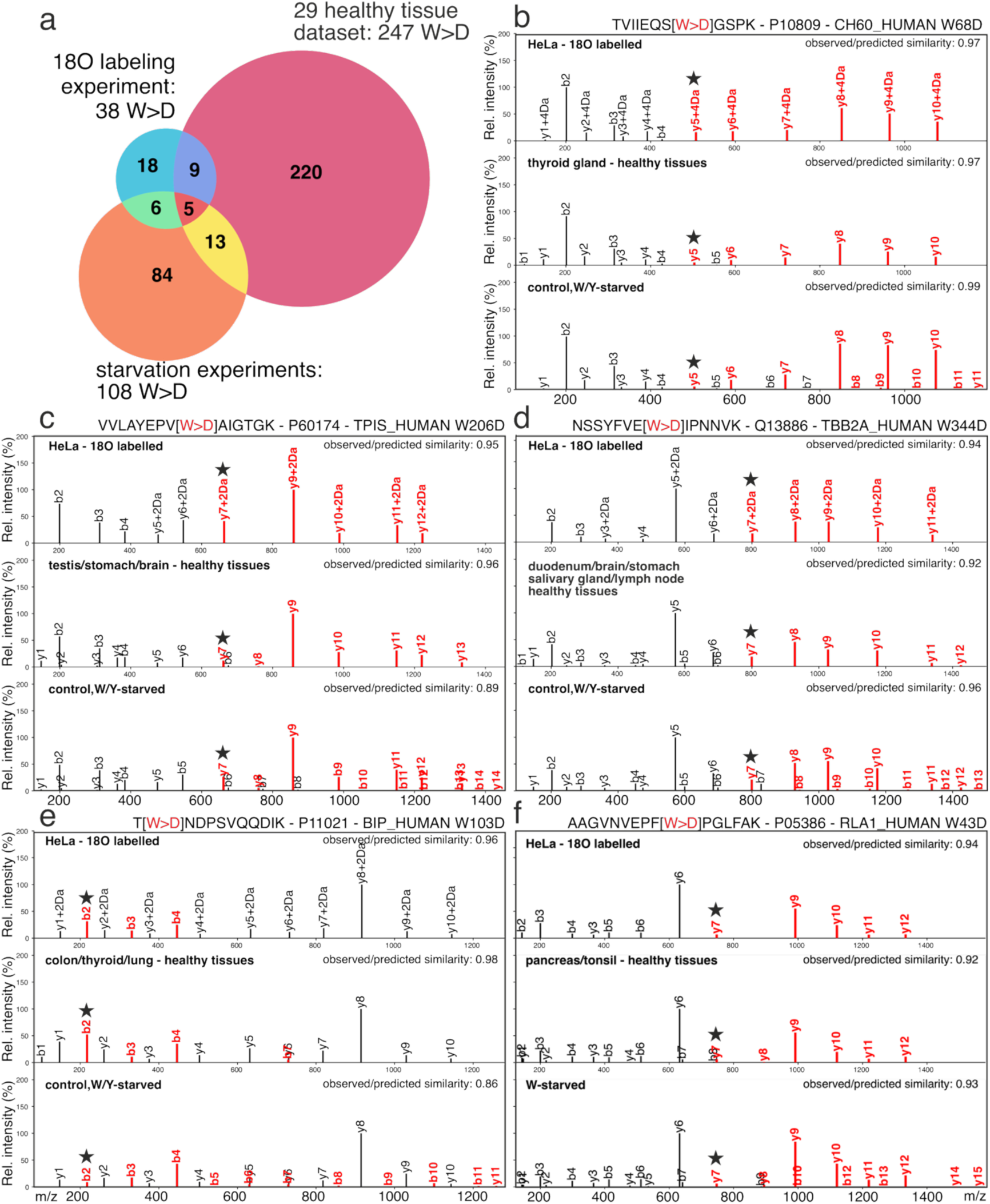
Analysis of W>D substitution-bearing peptides in three independent proteomic datasets. (a) Venn diagram showing the number of W>D identifications in each dataset, with overlaps indicating peptides detected in common. (b–f) Annotated MS/MS spectra of the five W>D peptides identified across all three datasets (¹⁸O-labeled HeLa cells, healthy tissues, and W/Y-starved human cell lines). Some W>D peptides were detected in multiple healthy tissues and starvation conditions (top left corner of each spectrum). Similarity of the experimental and predicted spectra in the top right corner of each spectrum. Fragments supporting the substitution are highlighted in red, the site-determining fragment is marked with a black star, and +2 Da / +4 Da suffixes on y-ion labels indicate the presence of one or two ¹⁸O atoms at the precursor C-terminus.

**Extended Data Fig. 9:**
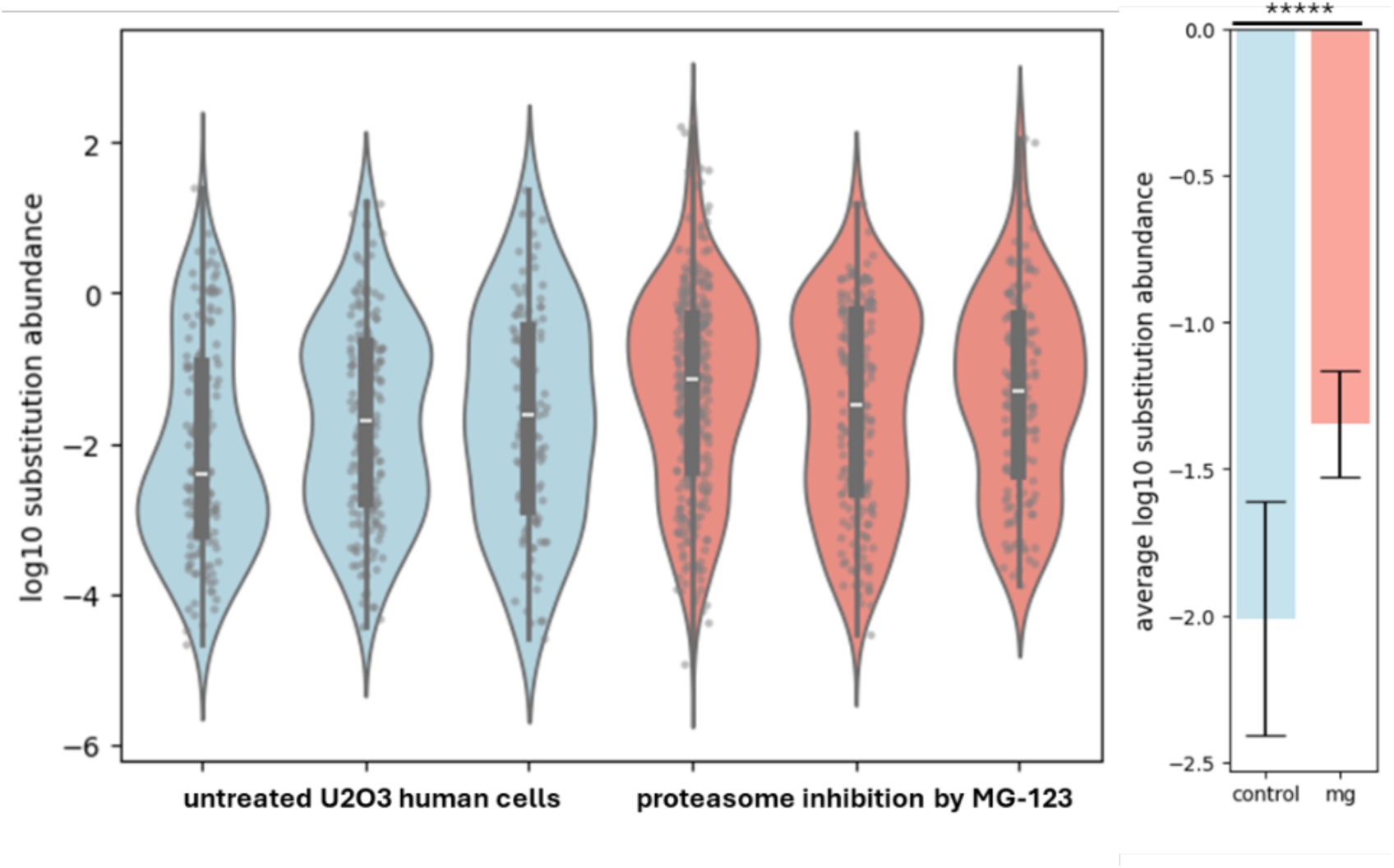
(Left) Violin plots showing the distributions of substitution abundances in untreated U2OS cells (blue) and MG-132–treated U2OS cells (red), shown separately for each biological replicate. (Right) Bar plots summarizing the significant differences between untreated and MG-132–treated conditions

**Extended Data Fig. 10:**
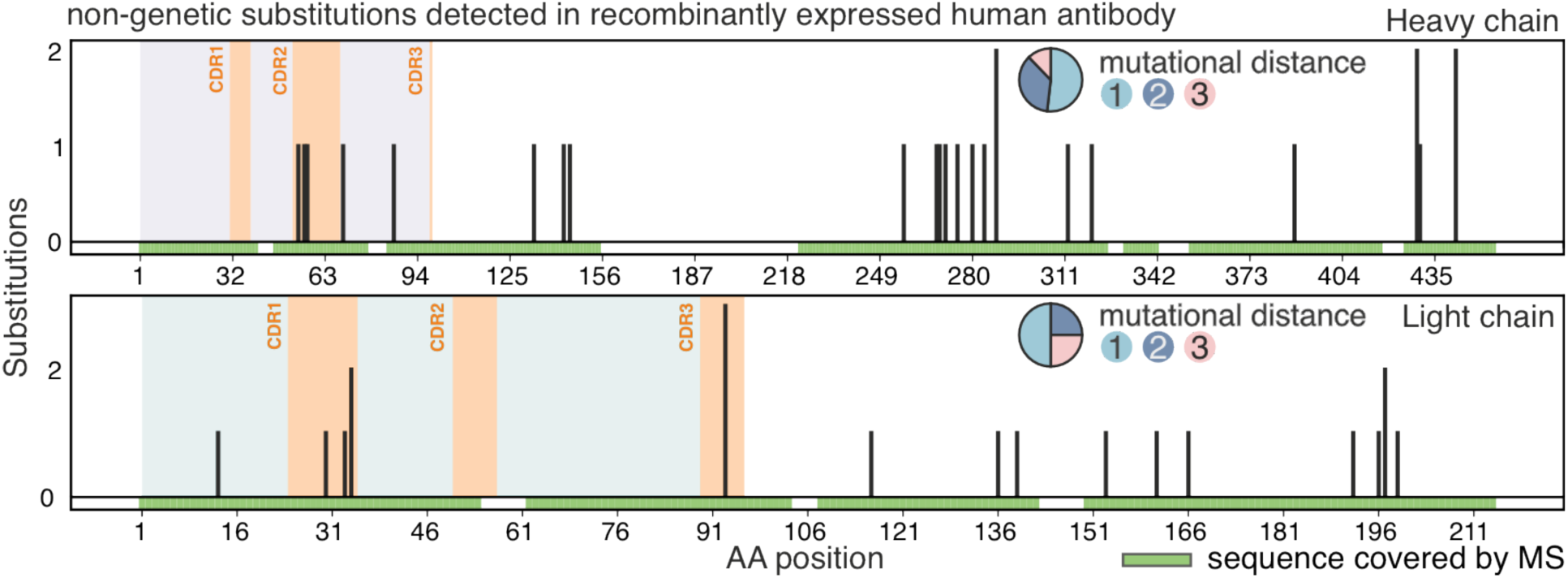
Substitutions detected in heavy (top) and light (bottom) chains of affinity purified human recombinant immunoglobulin protein R1. Amino acid positions covered by MS highlighted by the green panel above the x-axis. Each bar represents the count of different substitutions detected in respective positions of the protein. CDR areas highlighted in orange, framework regions in purple/green. Inset – proportions of the mutational distances among detected substitutions of each chain.

